# Temporal dynamics of noradrenaline release at fine spatial scales during motor learning

**DOI:** 10.1101/2025.10.29.685016

**Authors:** Xuming Yin, Nathaniel Jones, Aaron Jumarang, Tommaso Patriarchi, Simon X Chen

**Author notes:** These authors contributed equally (Xuming Yin, Nathaniel Jones).

## Abstract

Noradrenaline (NA) released from the locus coeruleus (LC) has been known to play pivotal roles in arousal, sensory processing, decision-making, and learning through global release across the entire neocortex. Recent studies have demonstrated heterogeneous and modular NA release in distinct brain regions and highlighted its spatiotemporal dynamics across the neocortex. However, the spatiotemporal specificity of NA release at fine scales within a single brain region remains unclear, and it has not been reported whether the release patterns evolve functionally throughout prolonged learning processes such as motor skill acquisition. Here, by employing *in vivo* two-photon imaging with various genetically encoded NA sensors, we reveal that behavior-induced NA release in the primary motor cortex (M1) is spatially heterogeneous at the scale of local microcircuitry. Over the course of learning, the release pattern is locally refined, achieving consistent spatial precision within M1. Intriguingly, pharmacological manipulations that disrupt the spatial specificity also alter local neurons’ activity and representations. Furthermore, LC-NA axonal calcium imaging uncovered two distinct temporal activity patterns, in which non-behavior-related ‘rapid’ axonal activity (sub-second duration events) profoundly affect the temporally precise behavior-induced ‘persistent’ axonal activity (seconds duration events). Closed-loop optogenetic manipulations that bi-directionally modulate non-behavior-related rapid events directly impacted the learning process. Together, our results provide novel insights into the temporal dynamics of NA release at fine spatial scales within one brain region and underscore the critical role of local NA specificity in regulating circuit plasticity during motor skill acquisition.

## INTRODUCTION

Noradrenaline (NA) is a neuromodulator widely distributed in the mammalian brain and is canonically associated with broad state modulation such as attention and arousal^1,2^. Over the past decades, accumulating evidence has also demonstrated its role in selective and modular circuit control of sensory processing^3,4^, cognitive function, behavioral flexibility, and decision making^5, 6^. NA in the forebrain originates primarily from the noradrenergic neurons in the locus coeruleus (LC), known to possess highly branched and long-projecting axons that innervate most cortical regions^7.^ Recent studies using rabies or other viral tracing methods8, and electrophysiological recordings^9^ have unveiled a high degree of functional heterogeneity in LC connectivity with prefrontal cortex, amygdala, and motor-related areas. In particular, it has been reported that distinct spatiotemporal activation dynamics of modular LC axons in two separate cortical regions play diverse roles during a head-fixed go/no-go auditory detection task^10^. However, the local, fine-scale spatiotemporal dynamics of NA release within a single cortical region are yet to be fully explored. Further, it remains unclear how these dynamics and release patterns are related to and evolve with a prolonged learning process such as motor skill acquisition.

The LC-NA system has been implicated in motor challenges observed in children with autism spectrum disorder (ASD)^11^. One of the most prevalent genetic risk factors for ASD is the microdeletion of the human 16p11.2 locus, which results in the loss of 550 kilobases of genomic DNA and haploinsufficiency of 29 genes ^12^.Notably, the predominant phenotypic manifestation observed in ∼60% of individuals with 16p11.2 deletion is motor difficulties, characterized by challenges in motor skills, notable clumsiness, speech delays, and difficulties in attaining developmental milestones such as crawling, walking, or sitting^13^. Deletion of the syntenic region of the 16p11.2 locus in mice *(16p11*.*2*^+/−^) recapitulates ASD-associated symptoms such as hyperactivity^14^ and social impairments^15^. We have previously shown that *16p11*.*2*^+/−^ mice exhibit a dysfunctional LC-NA system that causes delayed motor learning^16^; hence, it is critical to examine the role of NA release in *16p11*.*2*^+/−^ mice and explore its impact on the motor system. ^+/−^

In this study, we utilized newly developed genetically encoded NA sensors with *in vivo* two-photon imaging to probe the dynamic release of NA in the primary motor cortex (M1) of awake mice during a head-fixed motor learning task. We found that NA levels robustly increased during task-related movement in wild-type (WT) mice and can accurately predict movement epochs when used to train a support vector machine (SVM) classifier. Intriguingly, the initial release patterns were spatially segregated within M1 and became refined as learning progressed, and the spatial specificity of NA release patterns was critical in modulating local neurons’ activity and representations in M1. However, *16p11*.*2*^*+/−*^ mice exhibited attenuated and unspecific NA release patterns in the early stages of learning. We also performed *in vivo* two-photon calcium (Ca^2+^) imaging of the LC-NA projections in M1 to uncover the temporal dynamics of LC-NA activity. We revealed two types of axonal events, in which non-behavior-related rapid axonal activity affects the temporal specificity of behavior-induced persistent axonal activity. Lastly, to simulate the dysfunctions observed in *16p11*.*2*^*+/−*^ mice, we employed pharmacological and closed-loop optogenetic stimulation approaches to manipulate the spatial and temporal aspects of NA release in M1, respectively, and both manipulations were sufficient to alter the outcomes of motor learning. Our findings unveil that the spatial and temporal specificities of NA release at the scale of local microcircuitry within M1 is critical for learning and showcase how aberrant changes of NA release in an ASD model can preclude behavioral flexibility and disrupt learning.

## RESULTS

### Behavior-induced NA release during REs is spatially dynamic and undergoes refinement during motor learning

To investigate the dynamics of learning-related NA release in M1, we trained mice on a bi-directional rotating-disk task (1 hour/session, 1 session/day, 12 sessions; **Fig 1A);** WT mice showed a significant improvement in total distance travelled and the number of running epochs (REs) as previously reported **(Fig. 1B)**^16^. In this task, mice learn to refine their posture **(SFig. 1A-G)** to control the bi-directional rotation to run smoothly and continuously **(SFig. 1H)**. We expressed the genetically encoded fluorescent NA sensor, GRAB-NE1m (GRAB_NE_)^17^ in M1 and performed *in vivo* two-photon imaging to monitor NA release during the process. We observed that the sensor was preferentially expressed in neurites rather than somata; hence, we conducted imaging of GRAB_NE_-expressing neurites (∼ 350 x 350µm field of view (FOV); ∼200µm from the pia). To quantify the fluorescence, we segmented the entire FOV into a non-overlapping ROI grid (7×7 grid, each ROI is ∼50 x 50µm, **Fig. 1C)** and extracted the fluorescent signal from each ROI. An increase in GRAB_NE_ fluorescence was observed in many ROIs during REs **(Fig. 1D-E)**. To validate the fidelity of the GRAB_NE_ sensor to NA, we imaged the sensor continuously as the animal awoke from anesthesia, since it is known that LC-NA neurons are silenced during anesthesia^18-20^. We observed a robust increase in fluorescence signals as the mice awoke **(SFig. 2A-C)**, which unequivocally demonstrates that GRAB_NE_ is reporting NA activity. Inactive ROIs (e.g. due to blood vessels or limited GRAB_NE_-expressing neuropil) were excluded from the analyses (1.86 ± 0.61 ROIs/mouse; **Methods)**.

**Figure 1.**
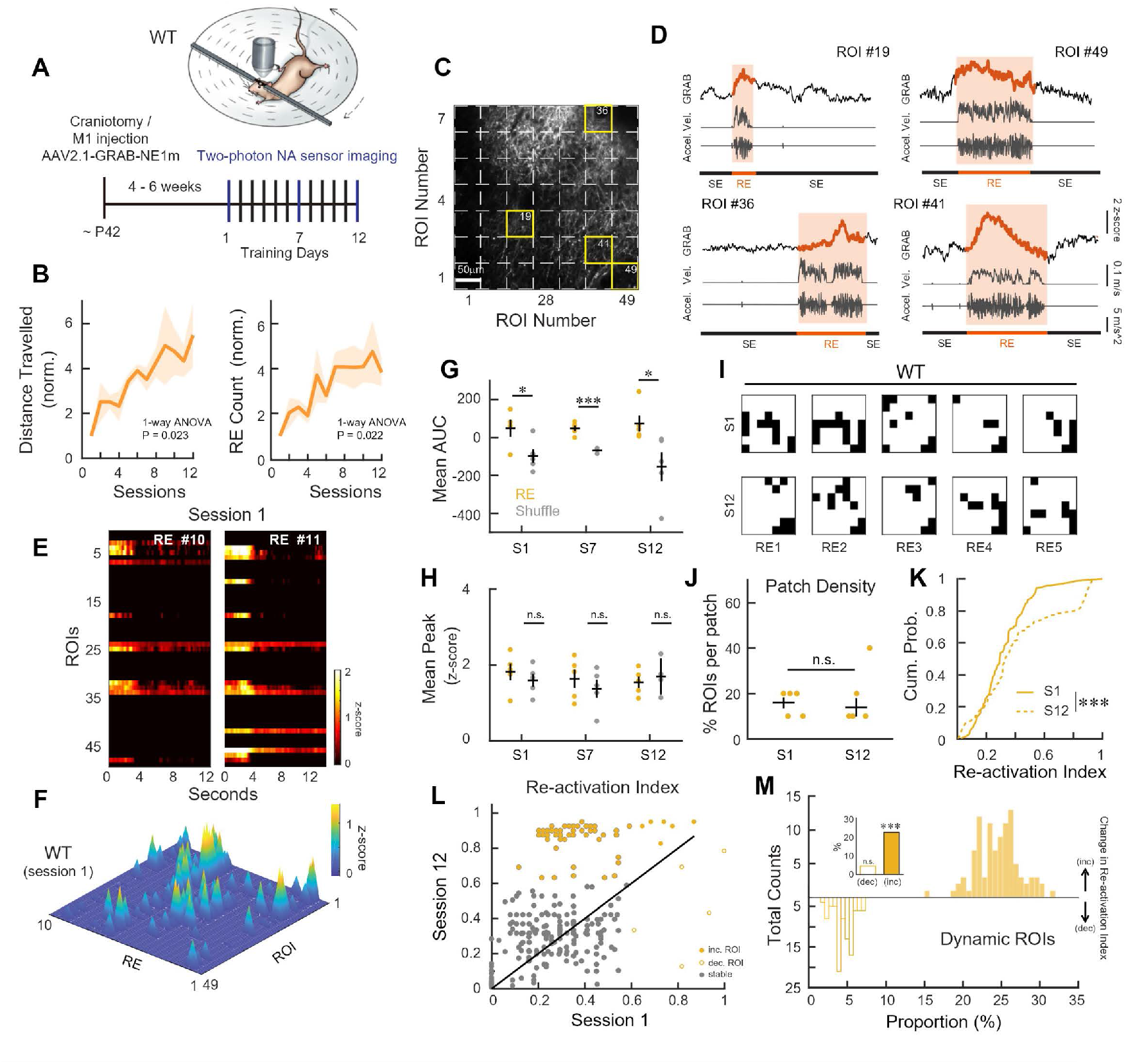
Behavior-induced NA release during REs is spatially dynamic and undergoes refinement during motor learning. **(A)** Top: schematic of bi-directional rotating-disk motor learning task. Solid and dashed arrows indicate the rotation direction of the rotating-disk. Bottom: Training and imaging timeline. **(B)** Total distance travelled (left) and running epoch (RE) counts (right) over 12 training sessions normalized to session 1. WT mice showed a significant improvement in both total distance travelled (F(11, 48) = 2.31, P = 0.023, one-way ANOVA; WT, N = 5 mice) and RE counts over training (F(11, 48) = 2.31, P = 0.022, one-way ANOVA). Solid line, mean value of each group; shading, s.e.m. **(C)** Example FOV containing ROI grid (7×7, white dashed line). Individual example ROIs in **D** are highlighted with yellow borders and numbered. Scale bar 50µm. **(D)** Example z-scored fluorescence traces during REs and the corresponding velocity and acceleration (RE: running epoch; SE: stationary epoch). Four selected ROIs from a WT mouse expressing GRAB_NE_ showed increases in fluorescence during REs. **(E)** Example heatmap of Z-scored fluorescence from two REs in session 1 from one WT mouse. ROIs do not show a global increase in GRAB_NE_ fluorescence across all ROIs during REs. **(F)** Surf plot illustrating average z-score from individual ROIs across 10 example REs in session 1 for one WT mouse. WT showed dynamic spatiotemporal NA release patterns in different REs. **(G)** Mean AUC during REs (yellow, N = 5 mice) for WT mice across learning compared to shuffled frame-matched SEs (grey). Two-sample T-test. REs show no change across learning: F(2,12) = 0.152, P = 0.86, repeated measures ANOVA. **(H)** Mean peak z-score values during REs (yellow) for WT mice across learning compared to shuffled frame-matched SE (grey). Two-sample T-test. **(I)** Active 7×7 ROI maps from one WT mouse; five running epochs in session 1 (top) and 12 (bottom). Black squares indicate active ROIs, white squares indicate inactive ROIs. **(J)** Mean patch density(# of ROIs in patch normalized to total active ROIs for each mouse) compared between sessions 1 and 12. One-tailed Wilcoxon rank sum test. **(K)** Cumulative probability of the re-activation index from individual active ROIs during REs (WT: n = 223 ROIs). Two-sample Kolmogorov-Smirnov test. **(L)** Re-ctivation index of all the active ROIs in session 1 (x-axis) plotted with session 12 re-activation index (y-axis) from all WT mice. Each circle is a ROI. Grey circles indicate ‘stable’ ROIs, filled circles indicate ‘increasing’ activity index, and empty circles indicate ‘decreasing’ re-activation index. Diagonal line indicates identity line (y=x). **(M)** Proportion of ROIs with increasing and decreasing activation indices from all WT mice. Filled histogram on top shows a 100x permuted distribution centered on proportion of increasing ROIs. Unfilled histogram on bottom shows a 100x permuted distribution centered on the proportion of decreasing ROIs. Inset shows proportions with significance compared to 5 percent (null; alpha level). Z-test. *P < 0.05, ***P < 0.001, n.s., not significant. Error bars indicate s.e.m.

**Figure 2.**
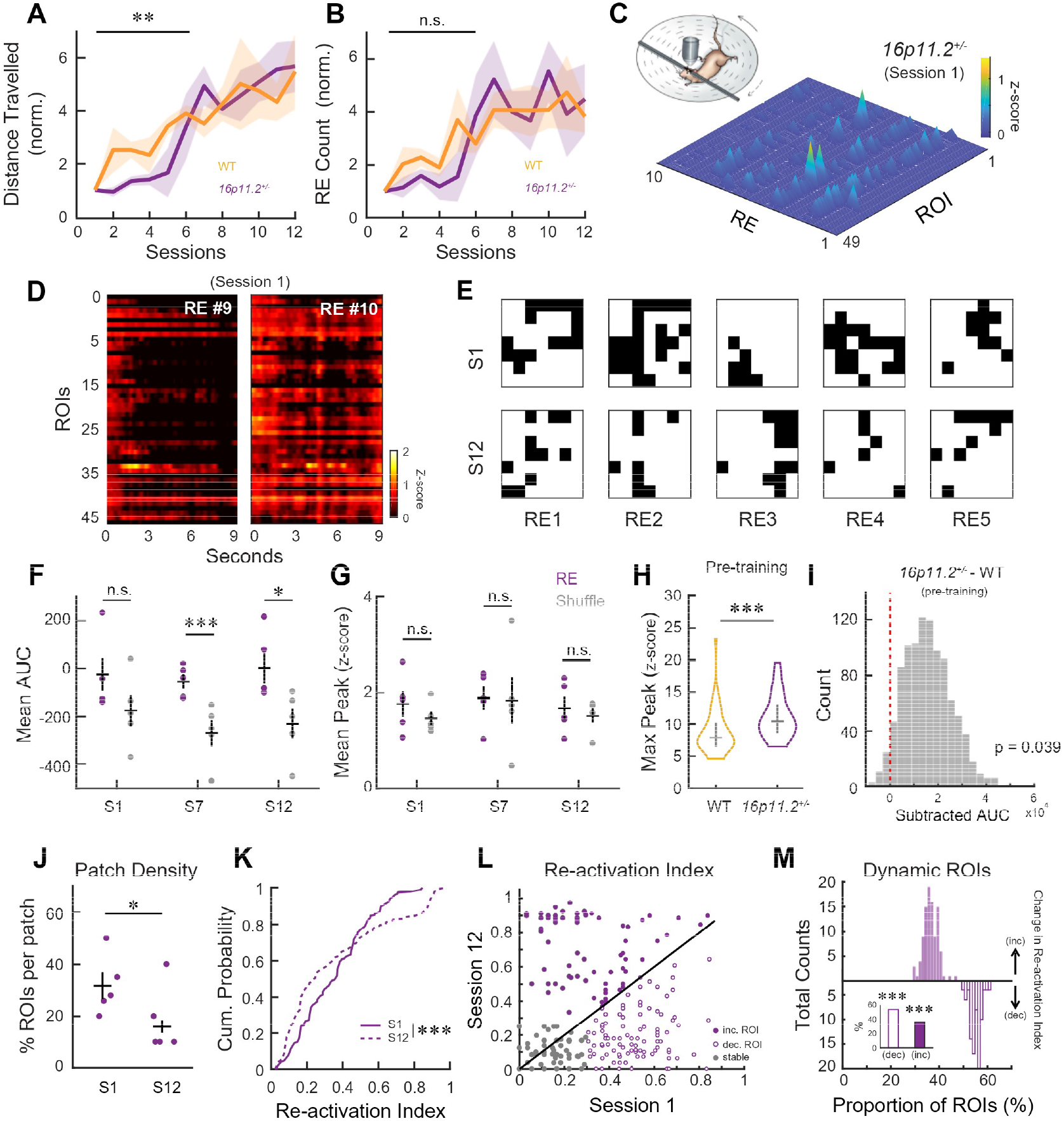
Behavior-induced NA release during REs in *16p11.2*^+/−^ mice is attenuated and spatially non-specific. **(A-B)** Total distance travelled **(A)** and RE counts **(B)** over 12 training sessions normalized to session 1. *16p11*.*2*^+/−^ mice showed a delay in improving distance travelled (Sessions 1–6:F(1,53) = 9.17, P = 0.004; WT, N = 5 mice; *16p11*.*2*^*+/−*^, N = 5 mice), but not in RE count (Sessions 1–6:F(1,53) = 1.74, P = 0.192). Solid line, mean value of each group; shading, s.e.m. **(C)** Surf plots of average z-score from individual ROIs across 10 example REs in session 1 for one *16p11*.*2*^*+/−*^ mouse, illustrating the release is weak and non-specific. **(D)** Example heatmap of Z-scored fluorescence from two REs in session 1 from one *16p11*.*2*^*+/−*^ mouse. Most ROIs in *16p11*.*2*^*+/−*^ mice showed attenuated and unspecific increase during REs. **(E)** 7×7 ROI maps from one *16p11*.*2*^*+/−*^ mouse; five running epochs in session 1 (top) and 12 (bottom). Black squares indicate active ROIs, white squares indicate inactive ROIs. **(F)** Mean AUC during REs *(16p11*.*2*^*+/−*^: purple, N = 5 mice) across learning compared to shuffled frame-matched SEs (grey). Two-sample T-test. REs show no change across learning: F(2,12) = 0.2507, P = 0.78, repeated measures ANOVA. **(G)** Mean peak z-score values during REs for *16p11*.*2*^*+/−*^ mice across learning compared to shuffled frame-matched SE (grey). Two-sample T-test. **(H)** Average maximum peak z-score of WT and *16p11*.*2*^*+/−*^ mice during the 10 minutes (1min per bin) pre-training baseline imaging. Mice were kept in a holding tube and did not run during this time. Two-tailed Wilcoxon rank sum test. **(I)** Histogram of differences in bootstrapped mean AUC between *16p11*.*2*^*+/−*^*and* WT mice from each bin of pre-training baseline. Red dotted line indicates expected difference under null hypothesis. Permutation test. **(J)** Mean patch density(# of ROIs in patch normalized to total active ROIs for each mouse) compared between sessions 1 and 12 in *16p11*.*2*^*+/−*^ mice. One-tailed Wilcoxon rank sum test. **(K)** Cumulative probability of the activation index from individual active ROIs *(16p11*.*2*^*+/−*^ mice: n = 242 ROIs). Two-sample Kolmogorov-Smirnov test. **(L)** Activation index of all the active ROIs in session 1 (x-axis) plotted with session 12 activation index (y-axis) from all the *16p11*.*2*^*+/−*^ mice. Each circle is an ROI. Grey circles indicate ‘stable’ ROIs, filled circles indicate ‘increasing’ activity index, and empty circles indicate ‘decreasing’ activity index. Diagonal line indicates identity line (y=x). **(M)** Proportion of increasing and decreasing activation indices, as in **1L**, for *16p11*.*2*^*+/−*^ mice. Z-test. *P < 0.05, **P < 0.01, ***P < 0.001, n.s., not significant. Error bars indicate s.e.m.

To comprehensively examine the spatial release pattern of NA throughout the entirety of learning, we tracked the same field of GRAB_NE_-expressing neuropil across session 1, 7, and 12 **(SFig. 3A-E)**. From the active ROIs, we first calculated the integrated area under the curve (AUC) of GRAB_NE_ fluorescence signal to examine if NA release was significantly increased during REs. We found an increase in the AUC during REs across all three imaging sessions in WT mice when compared to randomly selected stationary epochs (SEs) with matching duration **(Fig. 1G; Methods)**, but not in the amplitude of GRAB_NE_ fluorescence during REs **(Fig. 1H)**. This suggests that behavior-induced NA release during REs is consistent throughout learning. As a negative control, we imaged WT mice expressing GRAB_NEmut_ ^17^, which showed no change in fluorescence during REs at any stage of training **(SFig. 2F-I)**. Intriguingly, while we observed behavior-induced NA release during REs in M1, the release was not a global event across the entire FOV; instead, individual ROIs within M1 were activated in different REs **(Fig. 1E-F; SFig. 2J-K)**, forming ‘patches’ of multiple adjacent ROIs. We then examined the distribution of these NA release patches and investigated whether the size of the patches changed dynamically with learning. By constructing a binary mask of active ROIs, we defined a patch as a group of active ROIs that share one or more borders with other active ROIs **(Fig. 1I; Methods)**. On average, each RE contained ∼1.9 patches (composed of ∼4.6 active ROIs); however, the patch density did not change with learning **(Fig. 1J)**. Hence, we next calculated the re-activation index (percentage of REs in which a given ROI is active for each session) of individual ROIs during REs and SEs. We observed a population of ROIs that showed significantly higher re-activation indices by session 12 during REs compared to session 1 **(Fig. 1K)**, but not in SEs **(SFig. 3F)**, suggesting particular ROIs became more consistently re-activated specifically during REs after learning.

**Figure 3.**
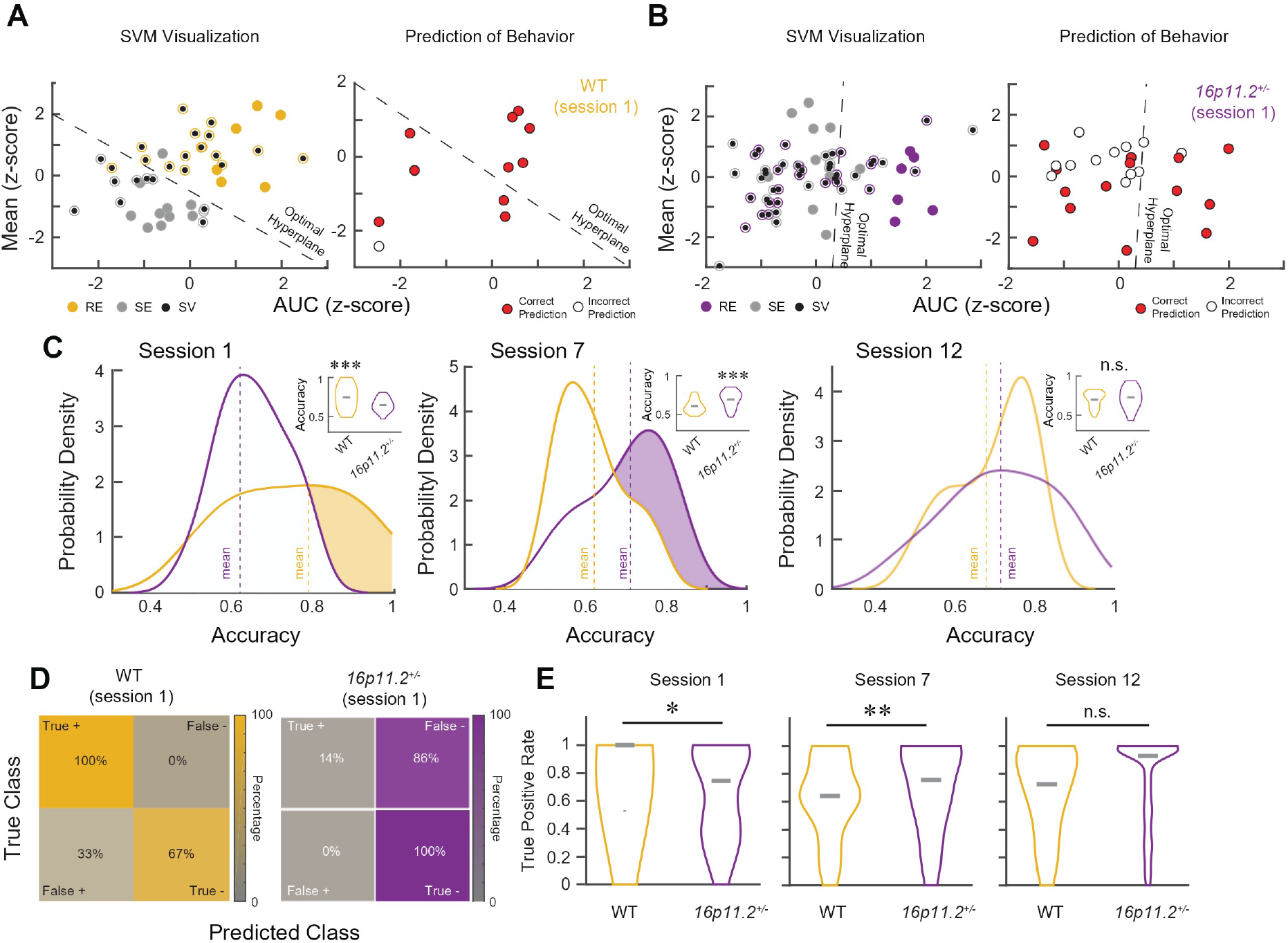
NA release signals are sufficient to predict animals’ behavior. **(A-B)** Representative two-dimensional# SVM visualization from one WT (A) and one *16p11*.*2*^*+/−*^mouse **(B)** in session 1. Left: REs (WT: yellow; *16p11*.*2*^*+/−*^: purple) and SEs (grey) from training dataset and putative* optimal hyperplane in relevant feature space. Black filled circles indicate epochs that are determined to be support vectors **(SV)** by the SVM classifier. Right: Hold-out epochs and respective classifier test predictions. Filled (red) circles indicate correct predictions, empty circles indicate incorrect predictions. # our SVM has four dimensions, one for each feature used to train * hyperplane shown is a 2D estimation for visualization purposes **(C)** Probability density of SVM prediction accuracy in session 1, 7, and 12, comparing WT and *16p11*.*2*^*+/−*^ mice. Shaded area indicates significantly higher accuracy. Vertical dotted lines indicate mean accuracy. Inset: violin plots indicate median of the same dataset. Wilcoxon rank sum test. Comparison to chance (0.5, i.e. 50%) level: session 1 (WT vs. *16p11*.*2*^*+/−*^, p<0.001), session 7 (p<0.001), session 12 (p<0.001). One-tailed T-test. **(D)** Example confusion matrices of a WT (left) and *16p11*.*2*^*+/−*^*mouse* (right) showing the percentage of True +, False+, True - and False - in session 1 (WT, n = 55 predictions; *16p11*.*2*^*+/−*^, n = 79 predictions). **(E)** Violin plots for prediction sensitivity (True + rate) for sessions 1, 7, and 12, comparing WT and *16p11*.*2*^*+/−*^ mice (N = 5 mice each). Grey bars indicate median. Wilcoxon rank sum test. *P < 0.05, **P < 0.01, ***P < 0.001, n.s., not significant. Error bars indicate s.e.m.

To identify the ROIs within this population, we tracked the re-activation index of each ROI from session 1 to 12. Since session 12 re-activation indices followed an approximately bi-modal distribution with a heavy tail and multiple local maxima **(SFig. 4H)**, we used a valley-based approach **(Methods)** to categorize each ROI with a threshold based on the local minima of the distributions, which we verified with a k-means clustering boundary. If an ROI was below the threshold, it was categorized as ‘basal’. Many ROIs remained basal throughout both sessions, so these ROIs were classified as stable (grey circles) as they did not change with learning **(Fig. 1L)**. To dissect the more dynamic ROIs (above the threshold), a linear separator (y=x, identity line) was employed to distinguish how many ROIs increased or decreased their activation indices from session 1 to 12 **(Methods)**. ROIs above (filled circles) the identity line were considered to have increased re-activation indices, while those below (open circles) were considered to have reduced re-activation indices **(Fig. 1L)**. We found that a significantly larger proportion of ROIs showed an increase in re-activation indices than a decrease **(Fig. 1M)**, indicating that a subpopulation of ROIs within M1 exhibited more spatially precise and consistent release of NA as motor learning progressed. We also confirmed that the emergence of spatial NA release was not due to changes in the sensor expression over time as mice trained only on session 1 and 12 did not show any increase in the re-activation index **(SFig. 3G-H)**. To further confirm our observations, we employed another recently developed NA sensor (nLightG; **SFig. 2D-E)**^21^. By applying the same imaging and analysis methods, an increase in AUC during REs was observed **(SFig. 4A-C)** as well as specific populations of ROIs becoming more consistently re-activated with learning **(SFig. 4D-E, J)**. Collectively, our results reveal heterogeneous NA release with patch-like patterns within M1, and the release becomes more consistent to achieve spatial precision as learning progressed.

**Figure 4.**
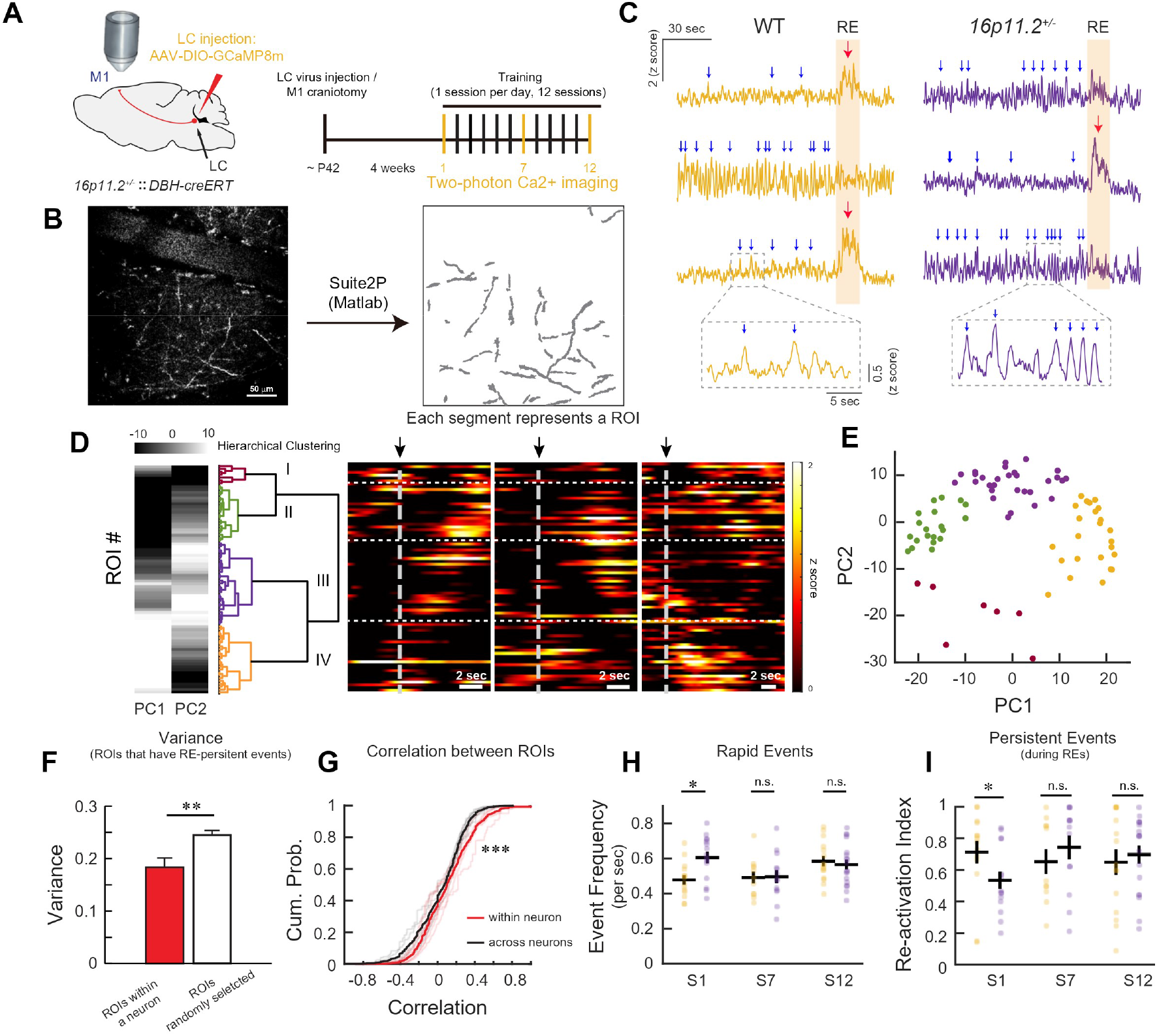
Excessive non-specific LC-NA rapid Ca^2+^ events and reduced behavior-induced persistent Ca^2+^ events during REs in *16p11.2*^+/−^ mice. **(A)** Left: schematic of LC-NA axonal Ca^2+^ imaging in M1 of *16p11*.*2*^*+/−*^::*DBH-CreERT* mice. Right: experimental timeline. Ca^2+^ imaging was conducted in session 1, 7 and 12. **(B)** Left: Example FOV containing GCaMP8m-expressing LC-NA axons in M1. Right: ROIs extracted by Suite2P. Scale bar, 50µm. **(C)** Example z-score normalized fluorescence traces of six ROIs from one WT mouse (left) and one *16p11*.*2*^*+/−*^*mouse* (right) in session 1. Blue arrows indicate ‘rapid’ Ca^2+^ events and red arrows indicate ‘persistent’ events. Shaded columns indicate RE. **(D)** Left: the first two principal components (PCs) (representing >80% of the variance) and hierarchical clustering dendrogram from one WT mouse. Right: example heat maps of the z-scored activity during three REs in session 1, showing axons in each neuron have distinct activity patterns during the onset of REs. Each row represents the activity of one axonal ROI. Black arrows and dashed grey lines indicate the onset of REs. Horizontal dashed white lines separate the clusters as determined by the hierarchical clustering (left). **(E)** PC distribution of each cluster from the example mouse in **(D)**. Each dot represents one ROI, and each color indicates one LC-NA neuron (cluster). **(F)** The variance in RE-persistent Ca^2+^ event occurrence among ROIs within a neuron was significantly lower than the randomly selected ROIs. Wilcoxon rank-sum test. **(G)** Correlation of RE-persistent Ca^2+^ event occurrence patterns within versus across neurons. ROIs within the same LC-NA neuron exhibited significantly higher similarity in the occurrence patterns **(SFig. 5C)** compared to ROIs in different neurons. Two-sample Kolmogorov-Smirnov test. **(H)** Rapid Ca^2+^ event frequency during SEs in session 1, 7 and 12 of WT and *16p11*.*2*^*+/−*^mice. *16p11*.*2*^*+/−*^ mice exhibited a higher frequency in session 1 (WT, n= 14 neurons from 7 mice; *16p11*.*2*^*+/−*^, n= 15 neurons from 5 mice) but not in later sessions. Each dot indicates one LC-NA neuron. Wilcoxon rank-sum test. **(I)** Re-activation index of persistent Ca^2+^ events during REs in session 1, 7, and 12. *16p11*.*2*^*+/−*^ mice showed reduced re-activation index in session 1 (WT, n = 14 neurons from 7 mice; *16p11*.*2*^*+/−*^, n = 15 neurons from 5 mice). Each dot indicates one LC-NA neuron. Wilcoxon rank-sum test. *P < 0.05, ***P < 0.001, n.s., not significant. Error bars indicate s.e.m.

We have previously reported that *16p11*.*2*^*+/−*^ mice exhibit delayed motor learning due to abnormal activation of NA neurons in the LC **(Fig. 2A-C)**^16^. Hence, we repeated NA sensor imaging in *16p11*.*2*^*+/−*^ mice to explore the NA release in M1 and its impact on motor learning. Strikingly, upon examining AUC and peak z-score of GRAB_NE_ fluorescence during REs in *16p11*.*2*^*+/−*^ mice, we did not observe a significant increase in AUC among the active ROIs in session 1 when compared to randomly selected duration-matched SEs **(Fig. 2F, G)**, suggesting mitigated NA release in REs among the active ROIs. However, the AUC became significantly higher in session 7 and 12 as training progressed **(Fig. 2F)**, and the time course of delayed NA release aligns with previously reported learning delay trajectories ^16^. Notably, the AUC and the peak z-score at the pre-learning baseline period in *16p11*.*2*^*+/−*^ mice was significantly higher than WT mice **(Fig. 2H-I)**, indicating that *16p11*.*2*^*+/−*^ mice may exhibit a higher basal level of NA that could hinder behavior-induced NA release during REs. Further investigation into the spatial release of NA in *16p11*.*2*^*+/−*^ mice also revealed a broad and unspecific release pattern in session 1, demonstrated by the significantly larger ROI patch size (∼ 2.56 patches per RE; ∼ 8.9 ROIs per patch) **(Fig. 2J)**. We again examined re-activation index of each ROI from session 1 and 12 in *16p11*.*2*^*+/−*^ mice as described above. While some ROIs in *16p11*.*2*^*+/−*^ mice exhibited increased re-activation indices in session 12 like WT mice, a larger proportion of ROIs in *16p11*.*2*^*+/−*^ mice exhibited a decrease in re-activation after learning **(Fig. 2K-M; SFig. 4I)**, demonstrating a more complex spatial reorganization of NA release in *16p11*.*2*^*+/−*^ mice.

These findings indicate that NA release in *16p11*.*2*^*+/−*^ mice is attenuated and non-specific during the initial phase of learning, and as training progresses, release patterns undergo a more complex refinement process wherein non-specific active ROIs reduce their activity, while a different population of behavior-related ROIs emerges to provide spatially consistent NA release. These observations support our previous hypothesis^16^ that low NA levels during REs contribute to the abnormal elevated activity among layer 2/3 excitatory neurons in M1 of *16p11*.*2*^*+/−*^ mice during the initial phase of learning. As training progresses and NA release increases, it may correct the abnormal activity in M1; however, the additional compensatory step results in delayed motor learning^16^.

### NA release predicts task-related movements better during the initial phase of learning

To further probe the relationship between NA release and task-related movements (REs), we used a support vector machine (SVM) algorithm and trained the classifier using GRAB_NE_ fluorescence features **(Fig. 3A-B; Methods)** from REs and SEs in both WT and *16p11*.*2*^*+/−*^ mice with a 20±10% hold-out rate. We then tested the trained classifier on held-out epochs to determine if NA release can accurately predict the behavior (RE vs. SE). To assess the training quality of the SVM, we calculated the hinge loss and found no significant difference between SVMs trained on WT and *16p11*.*2*^*+/−*^ mice (P = 0.8204; Two-sample t-test; hinge loss of 0.14 vs. 0.16 respectively), indicating reliable training. We next examined the overall accuracy of the classifier, and despite the similar training quality between the two groups, WT mice exhibited a higher prediction accuracy than *16p11*.*2*^*+/−*^ mice on session 1 **(Fig. 3C, left)**. This accuracy decreased in WT mice when the behavioral performance improvement plateaued (WT accuracy: S1: 74.6±2.0; S7 60.2±0.8; P < 0.001). Since LC-NA activity is related to attention and arousal^1,2^, it is possible that animals became less engaged in the task after they have mastered the movement. In contrast, *16p11*.*2*^*+/−*^ mice followed their characteristic delay, with lower accuracy in session 1 and an improvement in accuracy by session 7 (S1: 64.56±0.94; S7: 71.79±1.15; P < 0.05; **Fig. 3C, center)**. By session 12, once both groups had reached expert performance, their prediction accuracy was comparable **(Fig. 3C, right)**. We also investigated the decoding sensitivity of behavioral states (REs vs. SEs) by calculating the true positive rate (percentage of correctly predicted REs) across learning **(Fig. 3D)**. Similar to what was observed in the prediction accuracy, WT mice had higher decoding sensitivity in session 1, whereas *16p11*.*2*^*+/−*^ mice exhibited better decoding sensitivity in session 7 **(Fig. 3E)**, suggesting that the improvement of accuracy **(Fig. 3C)** is largely attributed to correct RE predictions. To confirm these predictive results, we trained an SVM classifier using the same fluorescent features from another NA sensor, nLightG, in WT mice that underwent motor learning. The fluorescent features of nLightG can also accurately predict REs, and there were no significant differences in prediction accuracy when compared to the SVM results from GRAB_NE_ across learning **(SFig. 4F-G)**, further supporting that behavior-induced NA release in REs is crucial for motor learning. Taken together, SVM analyses from both sensors utilized machine learning to confirm and expand the imaging results. By successfully decoding behavioral states using sensor features, the SVM analysis supports the claim that NA release is related to REs, and these sensor features contain sufficient information to distinguish between REs and SEs.

### Aberrant temporal activity patterns of LC-NA axons in *16p11*.*2*^*+/−*^ mice

Although GRAB_NE_ imaging has unveiled fine-scale spatial release patterns of NA in M1, it does not report the temporal activity patterns and heterogeneity of LC-NA neurons. To bridge this gap, we performed *in vivo* two-photon Ca^2+^ imaging of LC-NA projections in M1 to characterize the dynamic activity patterns of LC-NA axons in M1 during behavior. We injected AAV-DIO-GCaMP8m into the LC of *DBH-CreERT: :16p11*.*2*^*+/+*^(WT) and *DBH-CreERT::16p11*.*2*^*+/−*^ *(16p11*.*2*^*+/−*^) to express Ca^2+^indicator specifically in LC-NA neurons and waited four weeks following the surgery to image their axonal terminals in M1 **(Fig. 4A-C; SFig. 5A-B)**. We first used Suite2P to select ROIs in an unbiased manner and discarded the ones that did not meet the activity criteria **(Methods)**. We then performed principal component analysis (PCA) on the Ca^2+^activity of all axonal ROIs and used PCs for hierarchical clustering to classify ROIs into different LC-NA neurons (∼114 ROIs/mouse, ∼ 3 LC-NA neurons/mouse; **Fig. 4D-G; SFig. 5D)**^22^. Previous studies using two-photon Ca^2+^imaging of dopaminergic axons in the dorsal striatum have reported that Ca^2+^indicator fluorescent changes can be categorized into ‘rapid’ (sub-second duration events) and ‘persistent’ (seconds-duration events)^23^. Similarly, we observed rapid Ca^2+^ events (∼ 0.4 second length) and persistent Ca^2+^events (∼ 5 second length) among LC-NA axons in M1 (**Fig. 4C**). Surprisingly, the frequency of rapid Ca^2+^events in *16p11*.*2*^*+/−*^ mice during SEs was significantly higher than in WT mice in session 1 (**Fig. 4H)**, and these axons with abundant rapid Ca^2+^ events are also broadly dispersed across the FOV (**SFig. 5E-H**), resembling the elevated non-specific basal release of NA shown earlier **(Fig. 2H-I)**. In contrast, for the persistent Ca^2+^ events, we found that they occurred reliably during REs (71.3±7.2%) in WT mice, whereas the recurrence of persistent Ca^2+^ events during REs was significantly lower in session 1 of *16p11*.*2*^*+/−*^ mice **(Fig. 4I)**. To further compare axonal Ca^2+^ imaging with NA sensor imaging results, we applied the same 7 × 7 grid ROI method to the axonal images **(Methods)**. We found that subpopulations of the grid ROIs exhibited increased recurrence of persistent Ca^2+^ events during REs from session 1 to session 12 in both WT and *16p11*.*2*^*+/−*^ mice **(SFig. 5I-J)**, mirroring the learning-induced spatial refinement of NA release during REs from the NA sensor imaging. Given that sustained increases in action potential firing can generate prolonged axonal Ca^2+^ events lasting several seconds23, and that phasic firing of LC-NA neurons is well established to be task-related^1,3,24,25^, our results suggest that the lack of behavior-induced phasic firing of LC-NA neurons during REs in *16p11*.*2*^*+/−*^ mice may be a direct contributor to the delayed motor learning.

**Figure 5.**
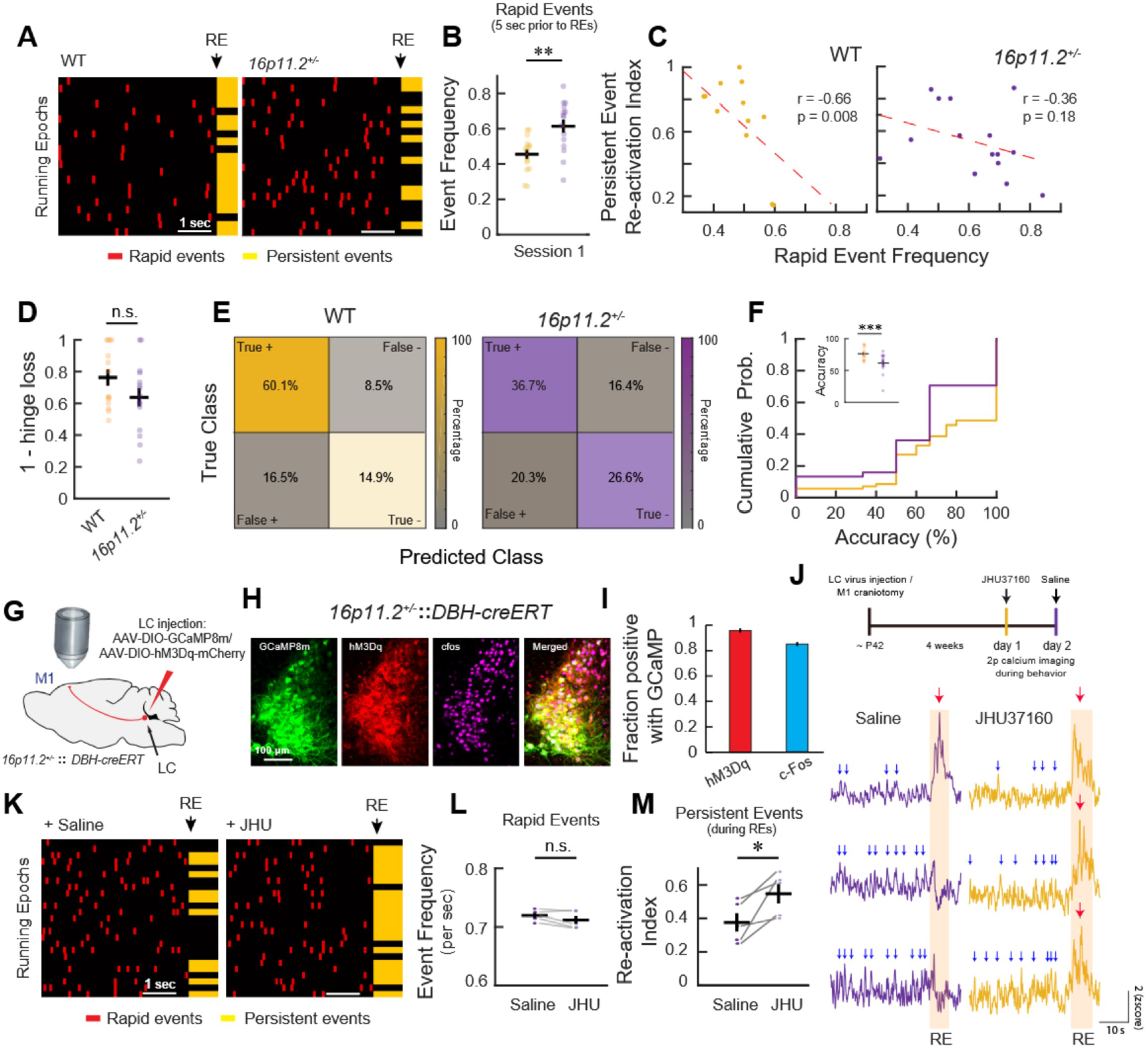
Temporally precise behavior-induced NA release during REs is critical for learning. **(A)** Representative plots showing the number of rapid Ca^2+^ events five seconds prior to the onset of REs in one WT (left) and one *16p11*.*2*^*+/−*^ mouse (right). Each row is 5 seconds leading up to a different RE from one neuron. Black arrow indicates the onset of REs. *16p11*.*2*^*+/−*^ mice exhibited fewer persistent Ca^2+^ events (yellow) and more rapid Ca^2+^ events (red) prior to the onset of REs. **(B)** Frequency of rapid Ca^2+^ events during the five second period prior to REs in session 1. *16p11*.*2*^*+/−*^ mice exhibited a significantly higher frequency of rapid Ca^2+^ events than WT mice. Each circle is a neuron. Wilcoxon rank-sum. **(C)** Correlation analysis between the persistent Ca^2+^ event re-activation index **(Fig. 4I)** and rapid Ca^2+^ events frequency **(Fig. 5B)** in session 1 of WT (left) and *16p11*.*2*^*+/−*^ mice (right). WT mice showed a negative correlation but *16p11*.*2*^*+/−*^ mice did not (Pearson R correlation coefficient, WT, r = −0.66, P = 0.01; *16p11*.*2*^*+/−*^, r=−0.36, P=0.18). **(D)** Hinge loss was comparable between WT and *16p11*.*2*^*+/−*^ mice in session 1. Wilcoxon rank-sum test. **(E)** Confusion matrix of SVM showing the True +, False +, True − and False − percentage in session 1 (WT, n = 188 predictions; *16p11*.*2*^*+/−*^, n = 177 predictions). **(F)** Cumulative probability of the prediction accuracy between WT and *16p11*.*2*^*+/−*^ mice in session 1. The insert plot shows mean accuracy from each LC-NA neuron. Wilcoxon rank-sum test. **(G)** Schematic of LC-NA axonal Ca^2+^ imaging in M1 of *16p11*.*2*^*+/−*^::*DBH-CreERT* mice with chemogenetic activation. **(H)** Representative images showing colocalization of GCaMP8m (green), hM3Dq-mCherry (red) and c-Fos (magenta) in LC-NA neurons. Scale bar, 100µm **(I)** Mean fraction of hM3Dq-mCherry cells and c-Fos-positive cells colocalized with GCaMP8m-positive LC-NA neurons (N = 5 mice). **(J)** Top: experimental timeline. Ca^2+^ imaging was conducted on day 1 with JHU37160 and day 2 with saline. Bottom: example z-scored fluorescence traces of three ROIs from one *16p11*.*2*^*+/−*^ mouse in the session with saline (left) and the session with JHU37160 (right). Blue arrows indicate ‘rapid’ Ca^2+^ events and red arrows indicate ‘persistent’ Ca^2+^ events. Shaded bars indicate RE. **(K)** Representative plots from one *16p11*.*2*^*+/−*^ mouse showing the number of rapid Ca^2+^ events five seconds prior to the onset of REs in the session with saline (left) and the session with JHU37160 (right). Each row is a RE. Black arrow indicates the onset of the RE. JHU37160 administration increased the occurrence of persistent Ca^2+^ events during REs while the rapid Ca^2+^ events prior to the onset of REs remained unchanged. **(L)** Chemogenetic activation of LC-NA neurons did not affect the rapid Ca^2+^ events during SEs (N = 5 mice). Paired t-test. **(M)** Chemogenetic activation of LC-NA neurons increased the re-activation index of persistent Ca^2+^ events during REs (N = 5 mice). Paired t-test. *P < 0.05, **P < 0.01, ***P < 0.001, n.s., not significant. Error bars indicate s.e.m.

Markedly, when we examined the relationship between the rapid and persistent Ca^2+^ events, we noticed that there were significantly more rapid Ca^2+^ events prior to RE onset in *16p11*.*2*^*+/−*^ mice **(Fig. 5A-B)**. Moreover, the recurrence of persistent Ca^2+^ events during REs was negatively correlated with the frequency of rapid Ca^2+^ events in WT mice but not in *16p11*.*2*^*+/−*^ mice **(Fig. 5C)**. To further investigate this relationship, we trained an SVM classifier using axonal Ca^2+^ activity features (5 seconds prior to each RE) to predict the occurrence of persistent events during REs **(Methods)**. SVMs trained on the activity features from both WT and *16p11*.*2*^*+/−*^ mice exhibit similar low hinge loss **(Fig. 5D)**, indicating reliable training from both groups. However, the prediction accuracy (correctly predicted the occurrence of persistent event) in WT mice was significantly better than *16p11*.*2*^*+/−*^ mice in session 1 (WT: 76.2±2.6%; *16p11*.*2*^*+/−*^: 61.5±3.9%; P < 0.001; **Fig. 5E-F)**. These observations align with the NA sensor results, in which excessive non-behavior-related NA release (from rapid axonal events) contribute to the elevated basal level of NA in *16p11*.*2*^*+/−*^ mice, thereby impeding temporally precise behavior-induced NA release (from persistent axonal events) during REs.

We have previously shown that chemogenetically activating LC-NA neurons was sufficient to rescue the delayed motor learning in *16p11*.*2*^*+/−*^ mice^16^. To understand how activation changes LC-NA axonal activity patterns in M1, we performed axonal imaging experiments while chemogenetically activating LC-NA neurons. We injected AAV-DIO-GCaMP8m and AAV-DIO-hM3Dq-mCherry into the LC of *DBH-CreERT: :16p11*.*2*^*+/−*^ to express Ca^2+^ indicator and the artificial excitatory receptor specifically in LC-NA neurons and waited four weeks following the surgery to image their axonal terminals in M1 **(Fig. 5G-I)**. Remarkably, chemogenetically activating LC-NA neurons enhanced the recurrence of behavior-induced persistent events during REs while having minimal impact on non-behavior-specific rapid events in *16p11*.*2*^*+/−*^ mice **(Fig. 5J-M)**, further highlighting that the temporal specificity of behavior-induced NA release during REs is critical for learning. We calculated an activity index for each

### Pharmacologically blocking NA reuptake abolishes the spatial specificity of behavior-induced NA release patterns during REs

Given the observed high basal levels of NA **(Fig. 2H-I)** and excessive non-behavior-related rapid axonal events in *16p11*.*2*^*+/−*^ mice **(Fig. 4H)**, we next asked whether pharmacologically increasing the basal level of NA in WT mice could affect behavior-induced NA release during REs and alter the learning process similar to *16p11*.*2*^*+/−*^ mice. To investigate this, we utilized the noradrenaline transporter (NET) inhibitor, atomoxetine, which has been shown to abolish NA re-uptake and prolong the presence of extracellular NA^26^. Using WT mice expressing GRAB_NE_ in M1, we performed *in vivo* two-photon imaging to monitor NA release for a 10min baseline session with the mouse in a holding tube, followed by intraperitoneal (i.p.) injection of atomoxetine (2mg/kg). We conducted another 30min imaging session while the mouse underwent training on the bi-directional rotating disk after the injection **(Fig. 6A)**, which permitted us to probe the effects of atomoxetine on behavior-induced NA release after blocking reuptake. We first evaluated NA levels between baseline and SEs and found that the peak z-score was increased during SEs after the administration of atomoxetine **(Fig. 6B, right)**. However, this was not observed in control mice **(Fig. 6B, left)**, indicating that blocking NA reuptake results in elevated basal levels of NA in M1, similar to the elevated baseline NA levels in *16p11*.*2*^*+/−*^ mice. Additionally, we found that the spatial release patterns of NA during REs in atomoxetine-treated mice appeared qualitatively broad and unspecific as in *16p11*.*2*^*+/−*^ mice **(Fig. 6C)**. We then examined whether elevated NA levels due to atomoxetine would perturb the behavior-induced NA release during REs. When quantifying the AUC during REs, compared to randomly selected duration-matched SEs, there was no behavior-induced increase in atomoxetine-treated mice **(Fig. 6D)**, although atomoxetine administration resulted in an increase in the patch size **(Fig. 6E)** and exhibited high re-activation indices compared to control mice **(Fig. 6F)**. These analyses demonstrate quantitatively that the release dynamics in atomoxetine-treated mice are broad and unspecific, reminiscent of NA release observed in *16p11*.*2*^*+/−*^ mice during REs **(Fig. 2)**.

**Figure 6.**
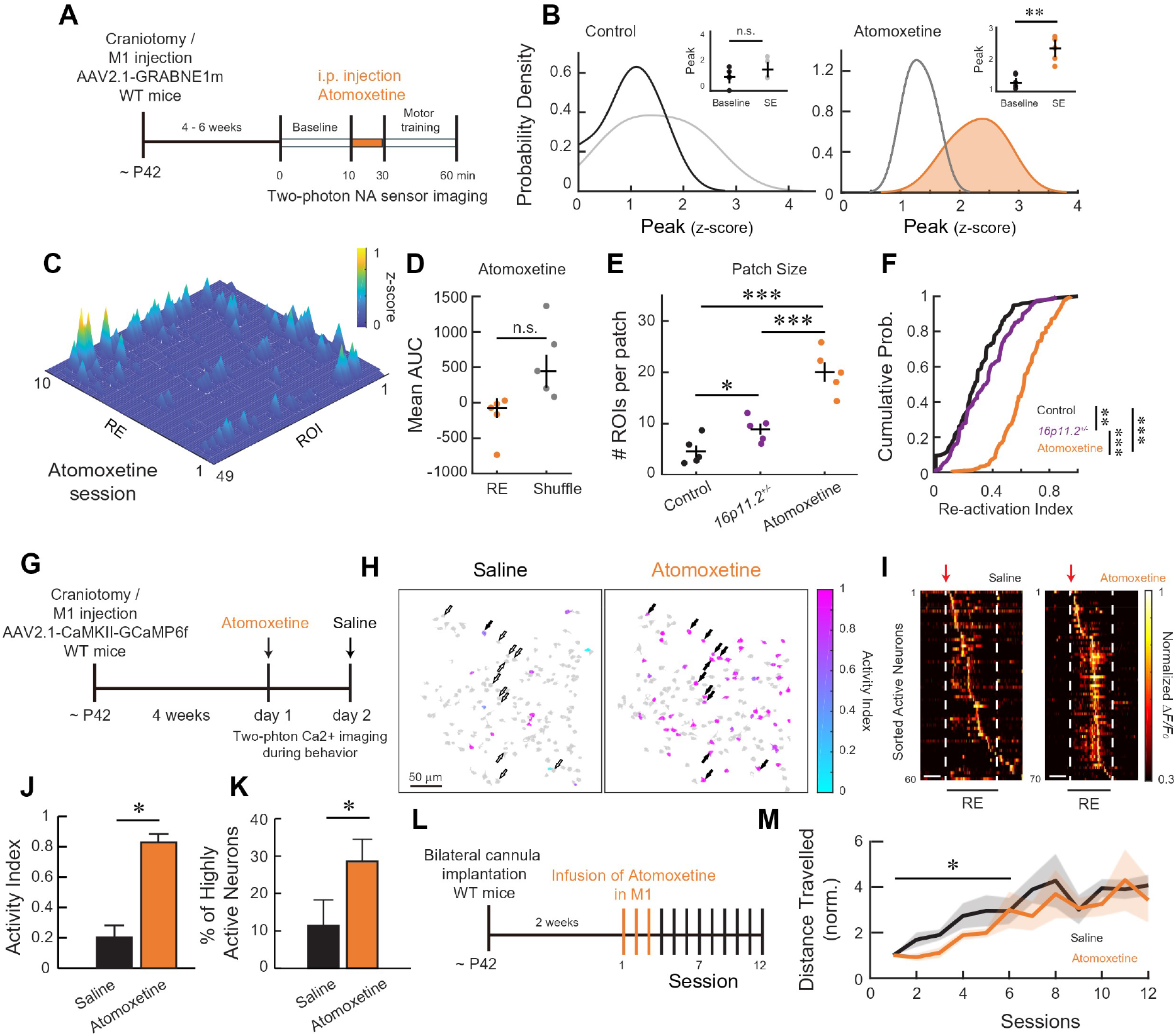
Pharmacologically blocking NA reuptake leads to non-specific NA release and altered neuronal representation in M1. **(A)** Experimental timeline. **(B)** Probability density of the peak z-score values from control and atomoxetine-treated mice during the baseline imaging period and SEs. While there was no change in control, mice treated with atomoxetine exhibited elevated NA levels during SEs, similar to *16p11*.*2*^*+/−*^ mice. Two-way ANOVA revealed a significant main effect of condition (F(1,4) = 6.43, P =0.0229), a significant effect of treatment (F(1,4) = 10.81, P = 0.005), and no significant interaction (F(1,4) = 0.78, P =0.3914). Inset: raw peak z-score data (each circle is a mouse). Post-hoc T-test with Bonferroni correction. Shading indicates significant difference from baseline. **(C)** Representative surf plot illustrating the average z-score from individual ROIs across 10 example REs in session 1 of one atomoxetine-treated mouse. The spatiotemporal release pattern resembles that of *16p11*.*2*^*+/−*^ mice **(Fig. 2C).** **(D)** Mean AUC during REs (orange) in atomoxetine-treated mice (N = 5 mice) compared to shuffled frame-matched SEs (grey). Atomoxetine-treated mice did not show behavior-induced NA release. One-tailed t-test. **(E)** Median number of ROIs per patch during REs (first session) in atomoxetine-treated mice compared to control mice and *16p11*.*2*^*+/−*^ mice (N = 5 mice in each group). Atomoxetine-treated mice exhibited larger patch size, aligned with *16p11*.*2*^*+/−*^ mice. Wilcoxon rank-sum test with Bonferroni correction. **(F)** Cumulative probability of the re-activation index from all active ROIs in WT, *16p11*.*2*^*+/−*^, and atomoxetine-treated mice. Atomoxetine-treated mice had more unspecific activation during REs, again similar to *16p11*.*2*^*+/−*^ mice. Wilcoxon rank-sum test with Bonferroni correction. **(G)** Experimental timeline of *in vivo* two-photon Ca^2+^ imaging of L2/3 excitatory neuron in M1 of WT mice. Repeated two-photon Ca^2+^ imaging of the same neurons was conducted on day 1 with atomoxetine and day 2 with saline injection. **(H)** Representative of excitatory neuron activity status across sessions from one mouse. Open arrows indicate the neurons that did not show high activity index; filled arrows indicate the neurons of high activity index. **(I)** Representative maximum-normalized activity of all active neurons from a single RE during the saline (left) and atomoxetine (right) sessions, aligned to RE onset (red arrow). Dotted white lines indicate the onset (red arrow) and end of the RE; scale bar, 2 s. **(J)** Atomoxetine administration increased neurons’ activity index during REs compared to the saline session (N = 4 mice). Paired t-test. **(K)** Atomoxetine administration increased the percentage of highly active neurons (N=4 mice) compared to the saline session. Paired t-test. **(L)** Experimental timeline with local atomoxetine infusion in M1 of WT mice. **(M)** Mean total distance travelled, normalized to session 1. Local inhibition of NA transporter in WT mice induced delayed motor learning (sessions 1–6: F(1,53) = 5.65, P = 0.021, two-way ANOVA). *P < 0.05, **P < 0.01, ***P < 0.001, n.s., not significant. Error bars and shading indicate s.e.m.

We next asked whether disrupting the spatial release patterns of NA would affect local neuronal activity and representations in M1. We injected AAV encoding Ca^2+^ indicator (GCaMP6f), driven by the CaMKII promoter (AAV-CaMKII-GCaMP6f), into M1 to express GCaMP6f selectively in excitatory neurons. Four weeks after surgery, we performed in *vivo* two-photon imaging and recorded the activity of hundreds of L2/3 excitatory neurons and followed the same population over 2 days (Day 1: atomoxetine ; Day 2: saline; **Fig. 6G)**. We calculated an activity index for each neuron, as described in our previous study (low positive index value indicates the neuron has a similar level of activity between REs and SEs, whereas a high positive index value indicates higher activity during REs than SEs; **Methods)**^16^. We found that when WT mice were injected with atomoxetine, more neurons showed increased activity during REs

(higher activity indices) compared to saline injection, demonstrating that changes in the basal NA level could alter the neuronal activity and representation in local excitatory neurons in M1 **(Fig. 6H-K)**. Lastly, to determine if atomoxetine treatment can impact the learning process, we implanted bilateral cannulae in M1 to specifically manipulate NA signaling locally. By infusing atomoxetine (3µg, 200nl) into M1 of WT mice 20 minutes prior to each training session during the early phase of learning (Session 1-3) to mimic the aberrant non-specific NA release observed in *16p11*.*2*^*+/−*^ mice, we found that WT mice exhibited delayed motor learning (F(1,53) = 5.65, P = 0.021, two-way ANOVA; **Fig. 6L-M)**, akin to the behavioral phenotype identified in *16p11*.*2*^*+/−*^ mice, without impacting animal’s general motor ability (**SFig. 6A-B**).

### Closed-loop manipulation of local LC-NA axon activity disrupts motor skill learning

While pharmacological manipulation has demonstrated the importance of spatially precise NA release, the impact of temporally precise NA release during task-related movements (REs) on motor learning remains to be determined. Based on the LC-NA axonal Ca^2+^ imaging results, we hypothesized that excessive non-behavior-specific rapid Ca^2+^ events preclude behavior-induced persistent Ca^2+^ events during REs and result in delayed motor learning. To test this hypothesis, we engineered a closed-loop optogenetic system, which tracks the running speed of the mouse while simultaneously employing DeepLabCut to monitor the pupil size (a proxy for LC-NA neuron activity) in real time and in an unbiased manner **(Fig. 7A, Methods)**. Previous research in both rodents and primates has established pupil size fluctuations as a dependable readout for LC-NA neuron activity^27-29^. In line with these findings, our results showed a notable increase in pupil size at the onset of REs, and the time course of pupil size and RE onset was more positively correlated than randomly selected epochs **(Fig. 7B-C)**. Pupil size during REs was also significantly larger than SEs **(Fig. 7C-D)**, as expected given the increased LC-NA activity during REs **(Fig. 4C, I**). To achieve targeted manipulation of LC-NA axonal activity in M1 in response to the mouse’s behavior, we first injected an AAV encoding cre-dependent channelrhodopsin-2 (AAV-DIO-ChR2-mCherry) into *DBH-CreERT* mice (ChR2 group) to specifically express ChR2 in the LC-NA neurons and implanted a glass window above M1 **(Fig. 7E; SFig. 6C-D)**. As a control, AAV**-** DIO-tdTomato was injected into a separate cohort of *DBH-CreERT* mice (tdTomato group). Using the closed-loop optogenetic system, we placed optic fibers above M1 and exclusively stimulated LC-NA ChR2-expressing axons (10-ms pulses, 10Hz, ∼20mW, 470nm) when both (1) animals were in SEs and (2) exhibited pupil size falling below a predetermined threshold (an indicator of low LC-NA neuron activity) **(Fig. 7F, Methods)**. Similar to the atomoxetine treatment, optogenetically activating LC-NA axons only during specific periods (SE + low pupil size) in the early phase of learning (session 1-3) was sufficient to induce delayed motor learning in WT mice (F(1,53) = 16.3, P = 0.0002, two-way ANOVA with post hoc Tukey’s test, **Fig. 7G**). Optogenetic stimulation in trained mice did not affect the animal’s motor ability nor their ability to initiate or terminate running **(SFig. 6E-H)**. Lastly, to further confirm the hypothesis that excessive non-behavior-related NA release (rapid events) in *16p11*.*2*^*+/−*^ mice could hinder behavior-induced NA release (persistent events) during REs, we repeated the closed-loop optogenetic experiments by silencing the LC-NA axons in M1 of *16p11*.*2*^*+/−*^ mice during specific periods (SE+ low pupil size). We injected AAV-DIO-ArchT-mCherry into *16p11*.*2*^*+/−*^*::DBH-CreERT* mice *(16p11*.*2*^*+/−*^ ArchT group) to specifically express the inhibitory opsin, ArchT, in the LC-NA neurons and implanted a glass window above M1 **(SFig. 6I)**. Following the same stimulation protocol (10-ms pulses, lOHz, ∼10mW, 590nm) and when the animal met the two conditions as described above, we found that silencing the excessive non-behavior-related rapid events in *16p11*.*2*^*+/−*^ mice rescued the delayed motor learning **(SFig. 6J)**, highlighting that bi-directional modulation of the temporal specificity of behavior-induced NA release can directly impact the learning process. Altogether, results from the pharmacological and closed-loop manipulations corroborate the hypothesis that the spatial and temporal specificity of NA release is necessary for effective motor learning. Moreover, non-specific release of NA in 16p11.2 deletion carriers can impinge upon NA dynamics that are crucial for learning or behavior engagement.

**Figure 7.**
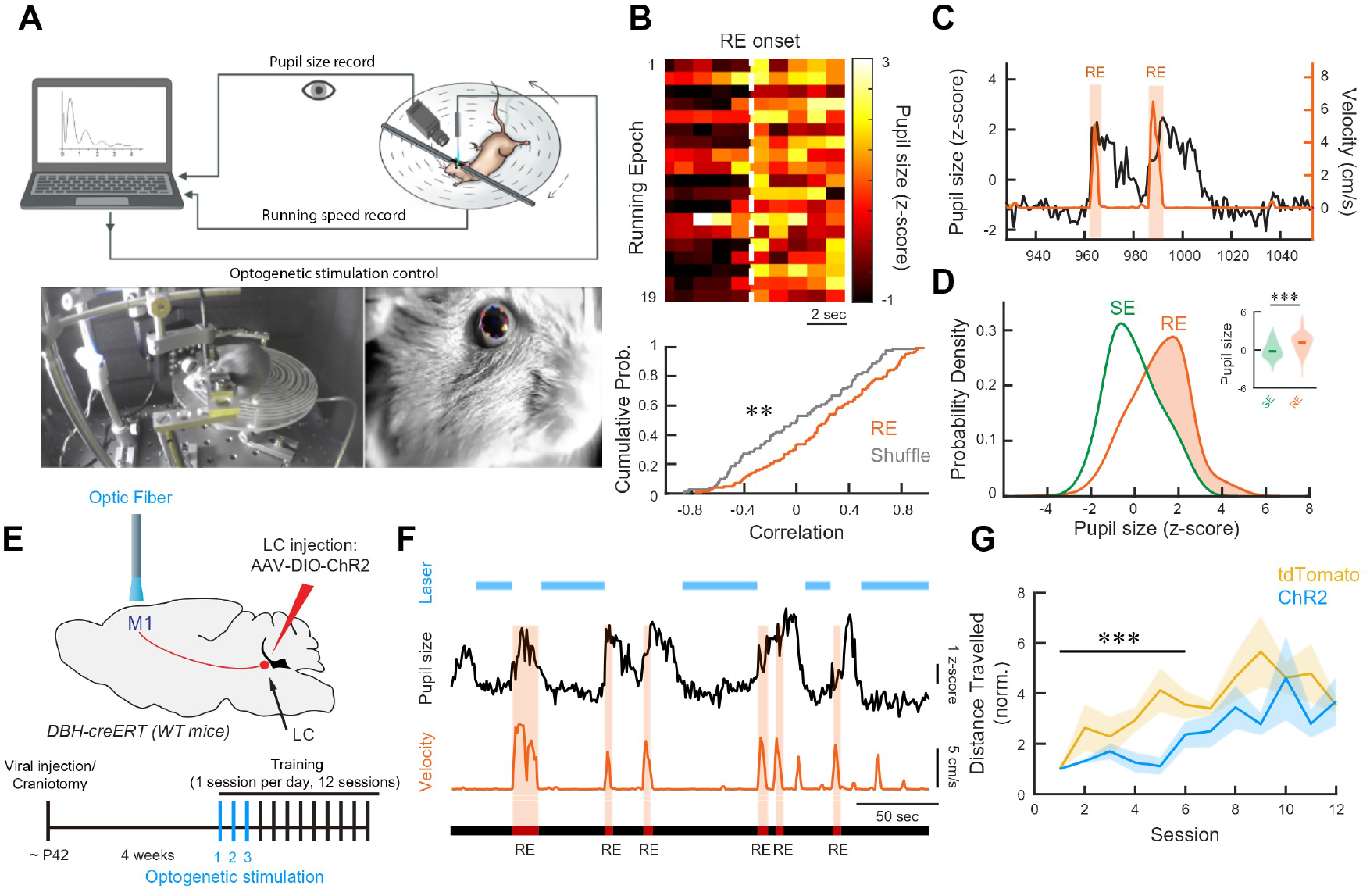
Closed-loop manipulation of LC-NA axonal activity by simulating non-specific Ca^2+^ is sufficient to alter the learning process. **(A)** Top, schematic of closed-loop manipulation of LC-NA axons in M1. Bottom left, experimental setup of closed-loop manipulation with one representative mouse performing the task. Bottom right, a representative image of online pupil size tracking using DeepLabCut. **(B)** Representative plot of the z-scored pupil sizes across all REs in session 1 of one WT mouse, aligned to the RE onset. Each row indicates one RE five seconds prior to the current RE. Dashed white line indicates the RE onset. Scale bar, 2 s. Bottom, cumulative probability of the correlations between the time course of pupil size change and binary running speed from all REs and randomly selected epochs of equivalent duration. **(C)** Representative plot from one WT mouse demonstrates that increase in pupil size coincides with RE onset. **(D)** Probability density of pupil size during REs and SEs. The insert violin plot shows the pupil size distribution and median. Pupil size in REs was significantly higher than SEs. Wilcoxon rank-sum test. **(E)** Schematic of the virus injection and optogenetic manipulation. Bottom, experimental timeline with optogenetic manipulation in WT mice. **(F)** Representative of closed-loop manipulation; LED was turned on based on pupil size and running velocity of one mouse in session 1. **(G)** Increasing non-specific LC-NA axonal activity during SEs induced delayed motor learning in WT mice (Sessions 1–6: F(1,53) = 16.3, P = 0.0002, two-way ANOVA; N = 5 mice). **P < 0.01, ***P < 0.001, n.s., not significant. Error bars and shading indicate s.e.m.

## DISCUSSION

The release patterns of NA were canonically considered to be global events due to the broad projection of LC-NA axons and the highly overlapping involvement of LC-NA neurons in various behavioral modalities^1,2,30^. However, a growing body of work from several groups have revealed heterogeneous and modular targeting of NA release across distinct brain regions^9,10,31-33^, shedding light on the spatiotemporal dynamics of LC-NA neurons and their projections. In the current study, we employed *in vivo* two-photon microscopy with NA sensor imaging and LC-NA axonal Ca^2+^ imaging in M1 and revealed distinct spatiotemporal release patterns of NA at the scale of microcircuitry within M1 during motor learning. Both GRAB_NE_ and nLightG sensor imaging demonstrated that NA release is reliably induced during task-related movements (REs) throughout learning. Moreover, machine learning classification using the fluorescent features from both NA sensors can accurately predict these movements, showcasing the importance of NA release in behavior. Intriguingly, the spatial distribution of NA release was heterogeneous within M1 and became refined as learning progressed, exhibiting more consistent NA release during REs. Furthermore, pharmacologically disrupting the spatial pattern of NA release alters the local neurons’ activity and representations in M1. Dopaminergic neurons have also been shown to exhibit isolated distal axonal activation locally in the striatum due to ectopic action potentials^34^. While it remains unclear which mechanisms trigger the ‘patch release’ of NA from LC-NA axons in M1, previous studies have proposed the ‘Glutamate Amplifies Noradrenergic Effects’ (GANE) model^24^, which suggests that local glutamate and NA can work mutually to enhance specific neuronal representations in the brain. It has also been shown that during motor learning, there is an emergence of reproducible L2/3 excitatory neurons activity patterns in M1^35^; hence, future work is essential to examine whether the spatial refinement of NA release may modulate local micro-circuitry by recruiting movement-specific neuronal ensembles during learning.

To better understand the temporal dynamics of NA release from LC-NA projections, we also performed *in vivo* two-photon Ca^2+^ imaging to record LC-NA axonal activity in M1 during motor learning. We observed two distinct types of Ca^2+^ events – ‘rapid’ (sub-second duration events) and ‘persistent’ (seconds-duration events), which have also been previously reported in dopaminergic axons in the striatum^23^. Axonal action potentials trigger the influx of Ca^2+^ into axons^36^, and a single action potential can lead to rapid Ca^2+^ events, whereas a long burst of action potentials results in sustained elevation of Ca^2+^ influx and gives rise to persistent Ca^2+^ events. Owing to the constraints of Ca^2+^ imaging, we are unable to verify whether rapid and persistent events correspond to the two classical LC-NA firing modes: ‘tonic mode’ − irregular but continuous baseline activity, and ‘phasic mode’ – bursts of higher-frequency activity during behavior^1, 37^. Despite this caveat, our observation of persistent Ca^2+^ events reliably occurring during REs in WT mice^27,38^ —together with behavior-induced NA release during REs from the NA sensor imaging—aligns with the ‘adaptive gain’ theory, wherein phasic activation of LC induces gain increases in downstream networks during task-related behavior^1^.

Intriguingly, we also revealed excessive rapid Ca^2+^ events during SEs and low occurrence of persistent Ca^2+^ events during REs in *16p11*.*2*^*+/−*^ mice. The Yerkes-Dodson ‘inverted U’ curve theory postulates that moderate tonic LC activity promotes optimal behavioral flexibility, while excessive or diminished tonic activity is associated with distractibility or lack of engagement respectively^39^. Moreover, it is known in the DA system that excessive extracellular DA can activate high-affinity presynaptic autoreceptors and lead to downregulation of axonal activity and attenuate the subsequent burst firing at the soma^34,40,41^. Hence, we hypothesized that the excessive rapid Ca^2+^ events during SEs could elevate the basal level of NA in *16p11*.*2*^*+/−*^ mice, thus lead to attenuated and unspecific behavior-induced NA release and alter the learning. To substantiate this hypothesis, we employed a closed-loop optogenetic stimulation system to precisely activate or silence non-behavior-related LC-NA axonal activity in M1 of WT mice or in *16p11*.*2*^*+/−*^ mice, respectively. By mimicking the non-specific LC-NA activity with the closed-loop optogenetic system, WT mice exhibited a delayed motor learning phenotype. In contrast, when the excessive LC-NA activity was silenced in *16p11*.*2*^*+/−*^ mice, the delayed motor learning was rescued.

The LC-NA system is complex and despite its well-regarded role in various behaviors, the specific mechanisms underlying its physiological action during learning are not yet clear. Our results demonstrate that behavior-induced NA release in M1 must be spatially and temporally precise during task-related behavioral states (REs), and disrupting this specificity can alter neuronal ensemble representations and affect the learning process. These findings will contribute to a growing field of work by characterizing the activity of LC-NA neurons and the pattern of NA release and highlighting the importance of NA signaling at fine scales within a single brain region during learning. Moreover, the observations in *16pl J*.*2*^*+/−*^ mice also align with studies on ASD children that showed increased tonic and reduced phasic activity of the LC-NA system compared to their typically developing peers^42^. Future work on dissecting the upstream factors that regulate LC-NA activity and its firing pattern in mouse models of ASD could pave new avenues for developing therapeutic strategies aimed at mitigating neural circuit dysfunctions associated with ASD.

## FIGURE LEGENDS

**SFigure 1.**
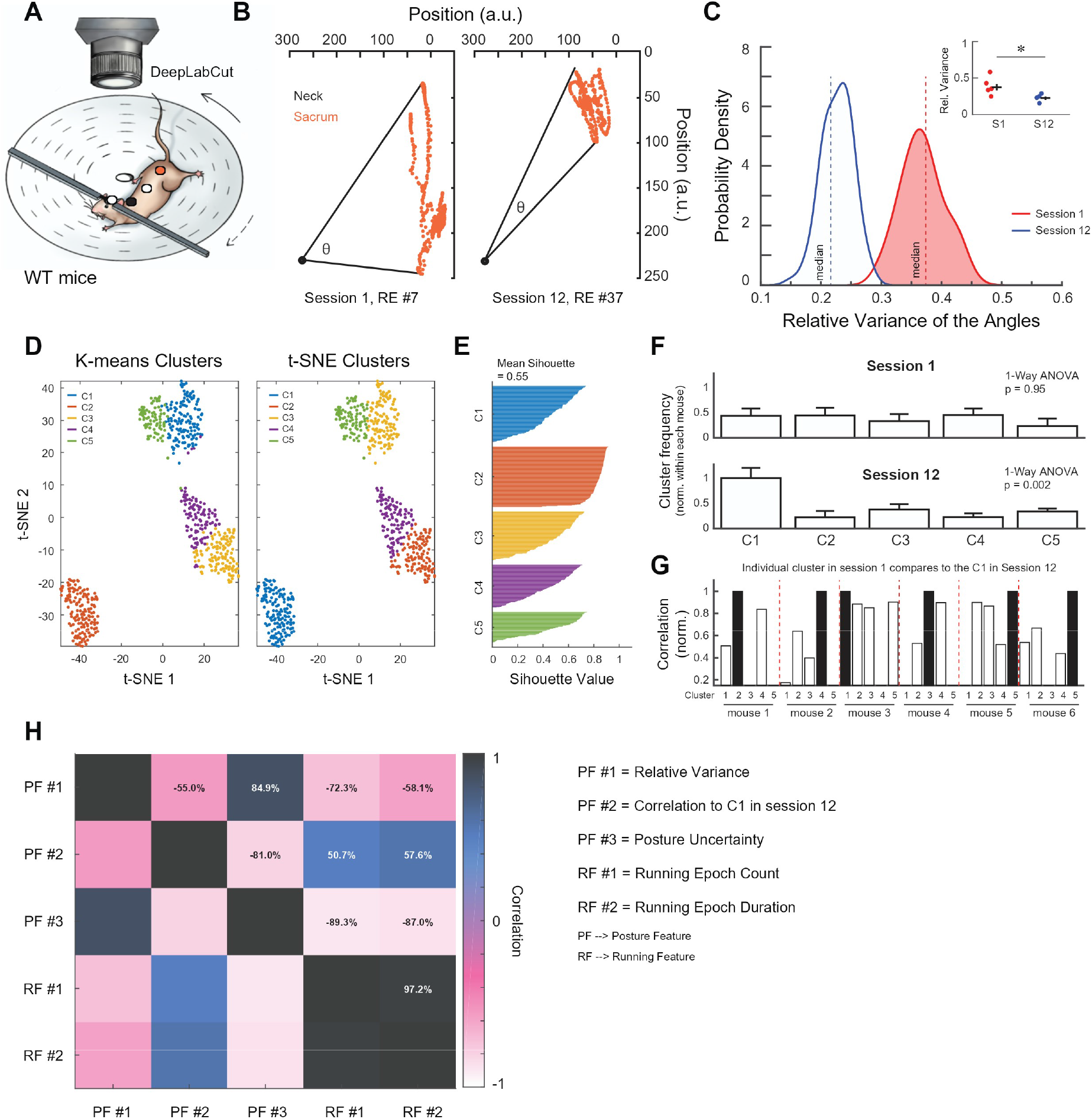
DeepLabCut analyses demonstrate postural refinement on the bi-directional rotating disk and link posture changes to running behavior. **(A)** Schematic of the bi-directional rotating-disk task recorded by a high speed and resolution camera from the top. DeepLabCut was used to unbiasedly track the position of head (white), neck (black), thorax (white), and sacrum (orange) across 12 sessions. **(B)** Representative plots of all the corporeal coordinates from the neck (black) and sacrum (orange) and the calculated angles from one RE of a mouse in session 1 (left) and session 12 (right). a.u., arbitrary unit. Mice posture became more consistent, and the angle variance was reduced with learning. **(C)** Probability density plot of mean bootstrapped (1,000x) relative variance of the angles in session 1 and 12 shows that the average relative variance was higher in session 1 than 12. Vertical lines indicate median. p<0.001. One-tailed T-test. Inset: mean relative variance of the angles from individual animals for session 1 and 12. p= 0.02. One-tailed T-test. **(D)** Example clustering from one mouse’s RE in session 1. Left: t-SNE projection of clusters obtained with k-means clustering. Right: k-means clustering applied to t-SNE transformed data. This comparison allows a visualization of the cluster consistency using dimensionality reduction. Cluster C1-5 indicate different postures. **(E)** Silhouette scores for the k-means clustering, which shows the cluster quality for each posture. Near 1 is very good, 0 is moderate, near -1 is poor. **(F)** Posture (cluster) frequency (in frames/posture) in session 1 (top) and 12 (bottom) normalized to first session. Postures in session 1 were more uniform and evenly distributed, indicating higher uncertainty (i.e. Shannon entropy). However, by the end of learning, posture frequency had shifted such that the primary posture was now preferred, resulting in lower uncertainty. P-values indicate the posture’s difference from one another. One-way ANOVA. **(G)** Correlation of each posture in session 1 to the preferred/primary posture (C1) in session 12. Black bars indicate the posture in session 1 that correlates the most with the eventual chosen posture in session 12. These postures become the primary postures from **F** (C1 in session 12). **(H)** Correlation matrix between postural features (PF) and running features (RF) across sessions 1, 4, 8, and 12. Correlations between the different feature types are highlighted with specific correlation values. Posture uncertainty (PF3) and RE count (RF1) were strongly negatively correlated, which suggests that mice have selected a specific posture permitting them to run more. Final preferred posture (PF2) and RE duration (RF2) were strongly positively correlated suggesting mice run longer as they refine to a preferred posture.

**SFigure 2.**
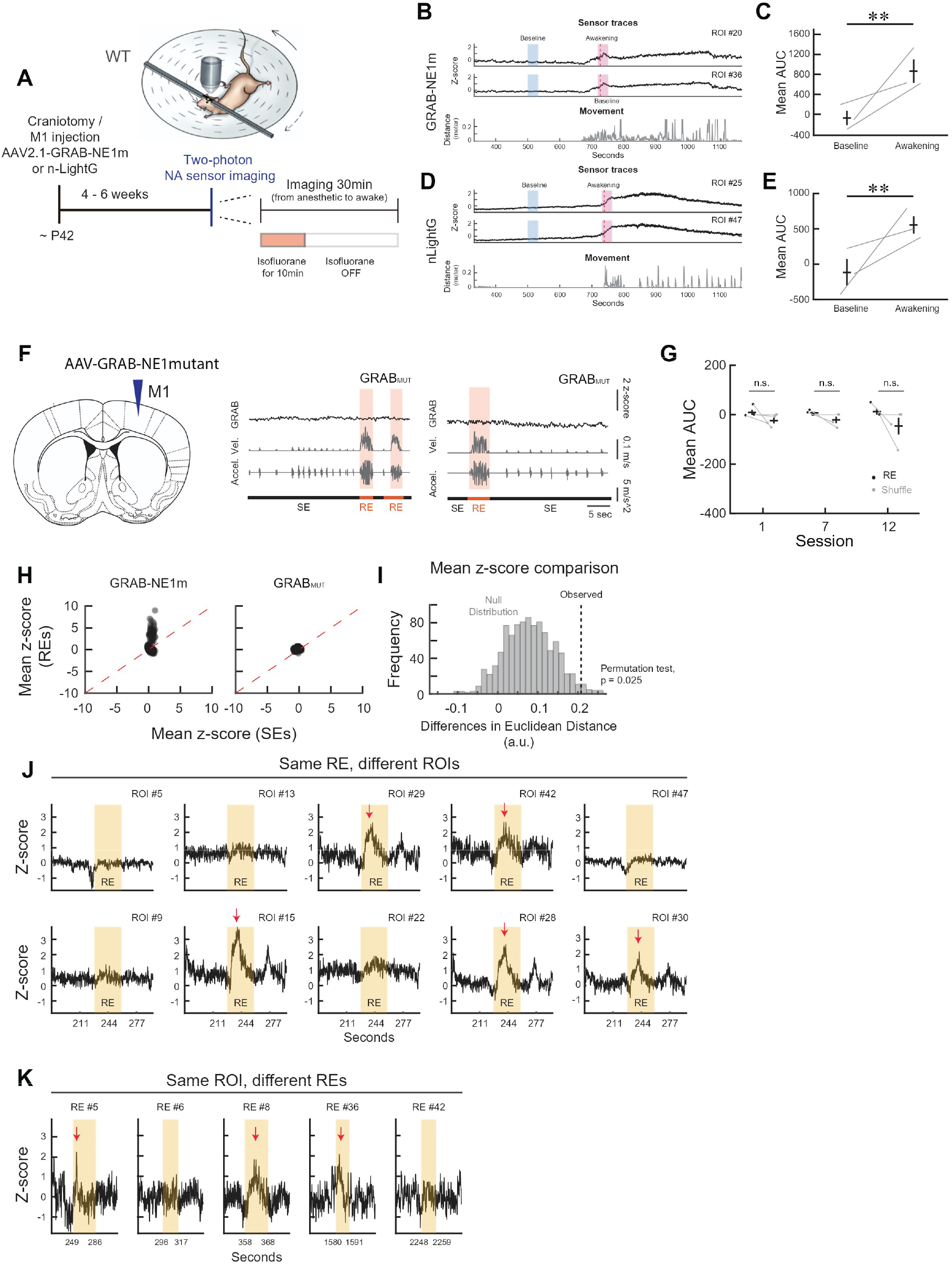
Validation experiments for GRAB_NE_ imaging of NA release. **(A)** Schematic and experimental design for isoflurane manipulation. **(B-C)** Example z-scored fluorescence traces (top) from two ROIs in one mouse anesthetized with isoflurane and the movement traces (bottom) **(B)**. Blue shaded region indicates window selected from anesthetized time period, pink shaded region indicates window selected during awakening. Red-dotted line indicates point of highest slope used to identify awakening. The mean AUC of the sensor fluorescence significantly increased as mice wakeup from anesthesia (**C)**. Two-sample T-test. **(D-E)** Same as **B-C** but for nLightG sensor. **(F)** Left: Schematic of virus injection. Right: Example z-scored fluorescence traces and the corresponding velocity and acceleration. Two selected ROIs from a WT mouse expressing GRAB_Nemut_. The mutant sensor showed no increase in fluorescence during REs. **(G)** Mean AUC during REs in GRAB_Nemut_ mice (N = 4 mice) compared to shuffled frame-matched Ses (grey). GRAB_Nemut_ mice did not show any behavior-inducedincrease in fluorescence throughout learning. Two-sample T-test. **(H)** Mean z-score values in GRAB_NE_ (left) and GRAB_Nemut_ (right) mice in REs (y-axis) plotted against the SEs (x-axis). GRAB_NE_ mice show dynamic increases in mean z-score during REs, whereas GRAB_NEmut_ mice show no dynamic activity in mean z-score during REs. **(I)** Difference in Euclidian distance between epochs (black dots in **H)** from GRAB_NE_ and GRAB_NEmut_ Black dashed line indicates observed value (true difference: GRAB_NE_ - GRAB_NEmut_). Grey histogram indicates simulated null distribution of differences. Observed difference is in the far tail of the null distribution, indicating that Euclidean distance of GRAB_NE_ epochs is larger than that GRAB_NEmut_, showcasing the larger dynamic range of the functional sensor. Permutation test. **(J)** Examples traces from 10 different ROIs in an RE from one mouse. **(K)** Example of different REs from one ROI in one mouse. *P < 0.05, ***P < 0.001, n.s., not significant. Error bars indicate s.e.m.

**SFigure 3.**
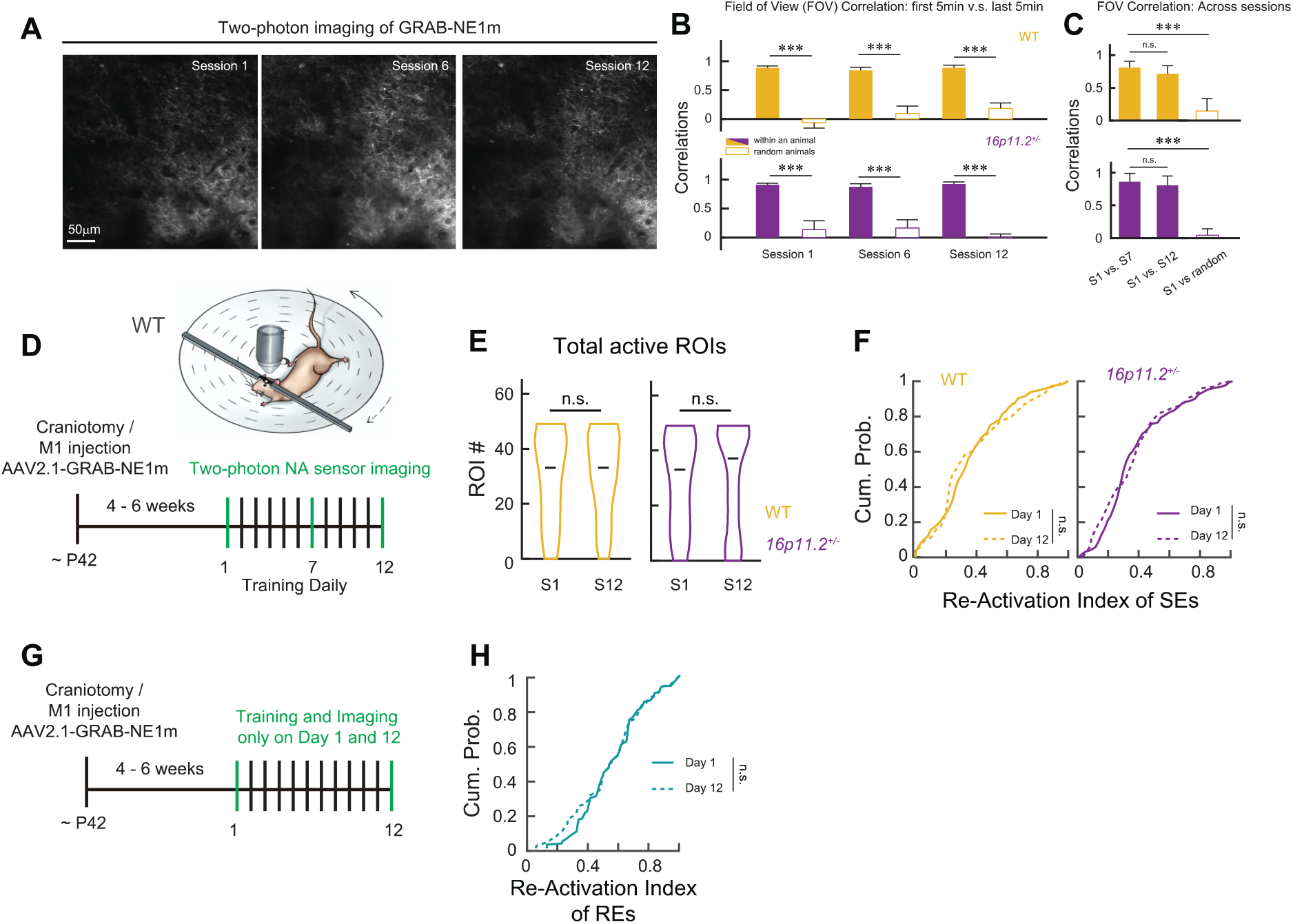
Validation of imaging FOV and ROI recruitment across sessions in trained and non-trained mice. **(A)** FOV (350 µm x µm) for an example mouse in session 1, 7, and 12. Scale bar is 50µm. **(B)** FOV pixel-wise correlations between first and last five minutes of an imaging session from the same mouse and randomly selected mouse for WT (top) and *16p11*.*2*^*+/−*^ (bottom). Two-tailed t-test. **(C)** Pixel-wise correlations from the same mouse and compared between session 1 and 7, session 1 and 12, and session 1 compared to another randomly selected mouse. Two-tailed t-test. **(D-F)** Schematic of the task and experimental timeline. Imaging occurred on days 1, 7, and 12 (same data from Fig 1 and 2). **(E)** Average number of active ROIs throughout an entire session for session 1 and 12 in WT (left) and *16p11*.*2*^*+/−*^ mice (right).Wilcoxon rank sum test. **(F)** Cumulative probability of the re-activation index from individual active ROIs during SEs in WT (left) and *16p11*.*2*^*+/−*^ mice (right). Wilcoxon rank sum test. **(G)** Experimental timeline. Imaging and training only occurred on days 1 and 12. **(H)** Cumulative probability of the re-activation index from individual active ROIs during REs on day 1 and day 12 in non-trained mice (N = 7 mice). Wilcoxon rank sum test.

**SFigure 4.**
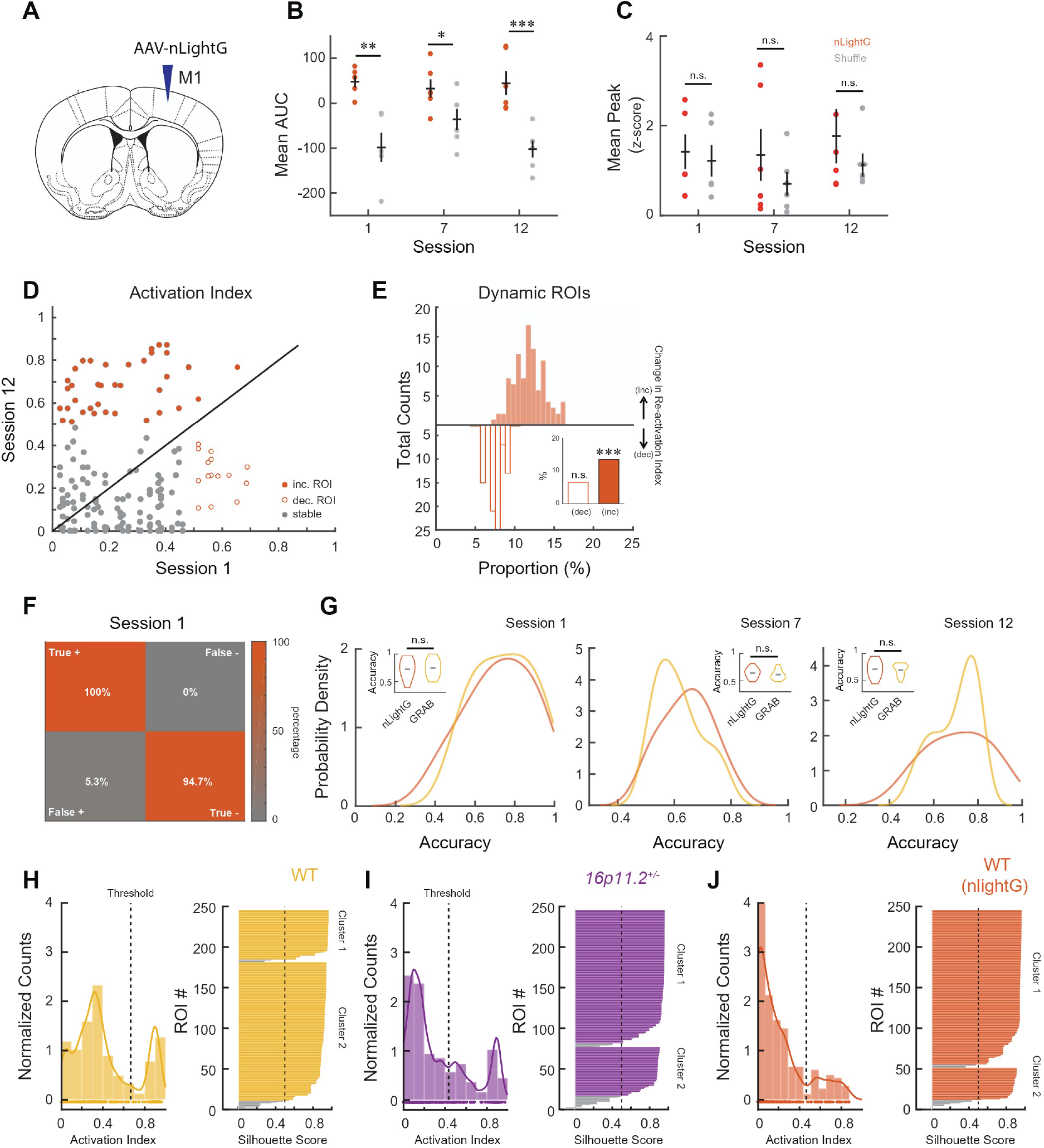
In vivo two-photon imaging of nLightG during learning across sessions. **(A)** Schematic of virus injection. **(B)** Mean AUC during REs (orange) in nLightG mice (N = 5 mice) compared to shuffled frame-matched SEs (grey). nLightG mice also showed an increase in fluorescence during REs, similar to what was observed in GRAB_NE_ imaging in **Fig. 1G**. Two-sample T-test. **(C)** Mean peak z-score values in nLightG mice did not change compared to shuffled frame-matched SEs (grey), similar to what was observed in GRAB_NE_ imaging in **Fig. 1H**. Two-sample T-test. **(D)** Re-activation index of all active ROIs in session 1 (x-axis) plotted as a function of session 12 re-activation index (y-axis) from WT mice. Each circle is a ROI. Grey circles indicate ‘stable’ ROIs, filled circles indicate ‘increasing’ reactivation index, and empty circles indicate ‘decreasing’ re-activation index. **(E)** Proportion of increasing and decreasing re-activation indices in nLightG mice as in **Fig. 1 & 2**. Z-test. **(F)** Confusion matrices nLightG mice showing the True +, False+, True − and False − percentage in session 1 (nLightG, n = 31 predictions). **(G)** Probability density of SVM prediction accuracy in session 1, 7, and 12, comparing nLightG and GRAB_NE_ mice. There were no differences in prediction accuracy between sensors. Inset, violin plot of median value. Wilcoxon rank-sum test. **(H-J)** Left: Distribution of re-activation index in session 12, including threshold to delineate ‘increasing’ or ‘decreasing’ ROIs. Right: Silhouette scores from k-means clustering of two clusters (i.e. increasing vs decreasing) for threshold validation in WT mice **(H)**, *16p11*.*2*^*+/−*^ **(I)**, and nLightG **(J)**.

**SFigure 5.**
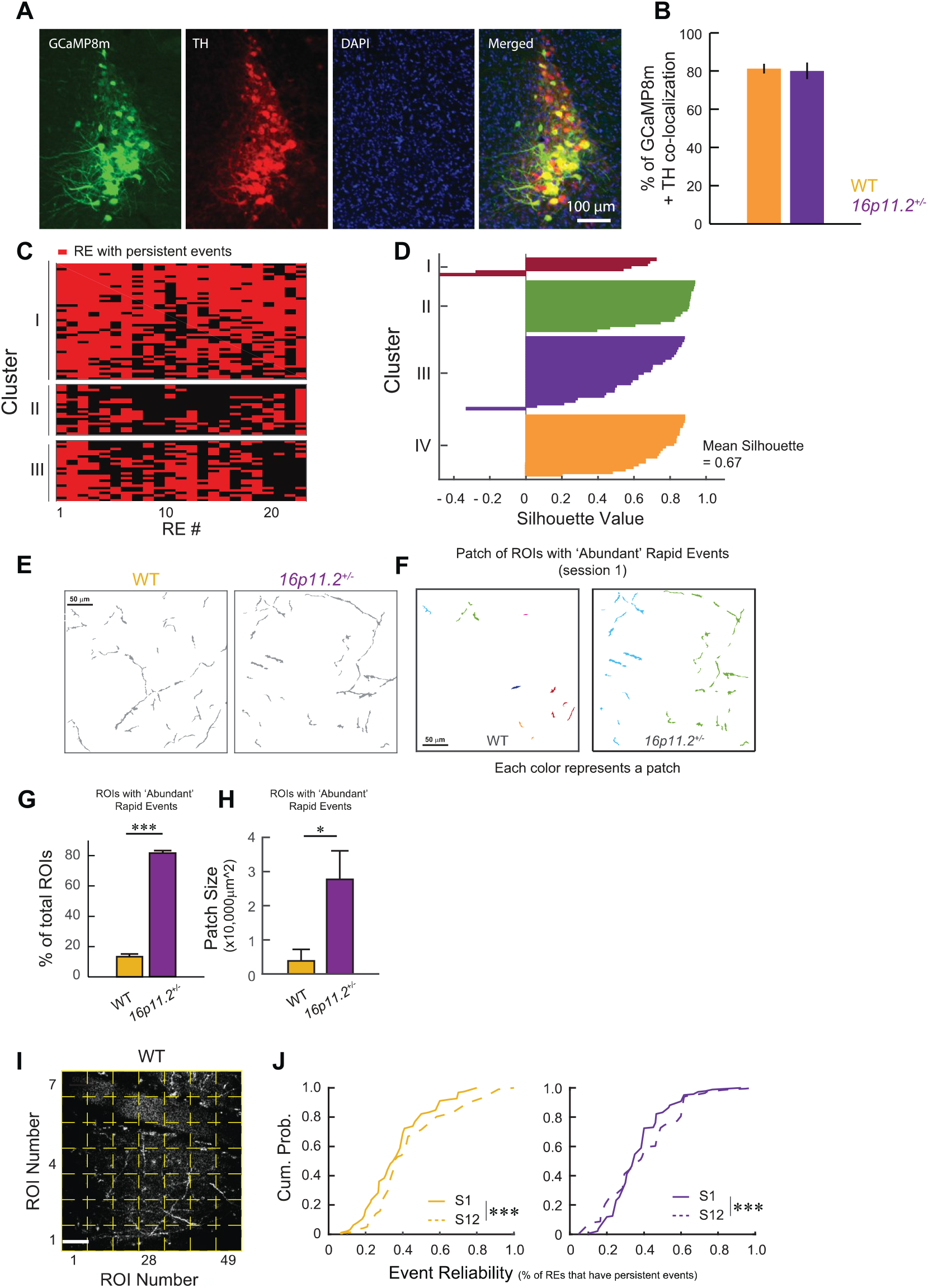
Validation of LC-NA axonal Ca^2+^ imaging. **(A)** Representative images showing GCaMP8m (green), TH immunofluorescence (red), DAPI (blue), and co-localization (merge). **(B)** Mean fraction of GCaMP8m-expressing cells co-localized with TH+ NA neurons (N = 5 mice) **(C)** Representative plots showing persistent Ca^2+^ event occurrence patterns during REs from one WT mouse. Each row corresponds to an ROI. ROIs from the same neuron exhibiting similar patterns of persistent event occurrence. **(D)** Silhouette scores for hierarchical clustering of a representative mouse (same mouse as in Fig. 4D–E), showing the clustering quality for each cluster (LC-NA neuron). Scores near 1 indicate well-clustered neurons, values around 0 suggest ambiguous assignments, and scores near −1 reflect poor clustering. **(E)** Representative image of ROIs extracted by Suite2P from one WT mouse (left) and one *16p11*.*2*^*+/−*^ mouse (right). Scale bar, 50 µm. **(F)** Representative image showing the physical distribution of LC-NA axons in the imaging FOV that had abundant rapid events in session 1 from one WT mouse (left) and one *16p11*.*2*^*+/−*^ mouse (right). Each segment represents a ROI, and each color indicates a distinct patch. **(G)** *16p11*.*2*^*+/−*^ *mice* had a significantly higher percentage of LC-NA axons that exhibited abundant rapid events than WT mice. Wilcoxon rank-sum test. **(H)** *16p11*.*2*^*+/−*^ *mice* showed larger patch size of LC-NA axons that exhibited abundant rapid events than WT mice. Wilcoxon rank-sum test. **(I)** Example FOV from axonal Ca^2+^ imaging of a WT mouse, and a 7×7 grid outlined by yellow dashed lines is applied to the FOV. The fluorescence from each grid is extracted as an ROI, similar to what was performed with the NA sensor imaging. Scale bar 50µm. **(J)** Cumulative probability of the re-activation index from individual active grid ROIs in the axonal Ca^2+^ imaging analysis. Two-sample Kolmogorov-Smirnov test.

**SFigure 6.**
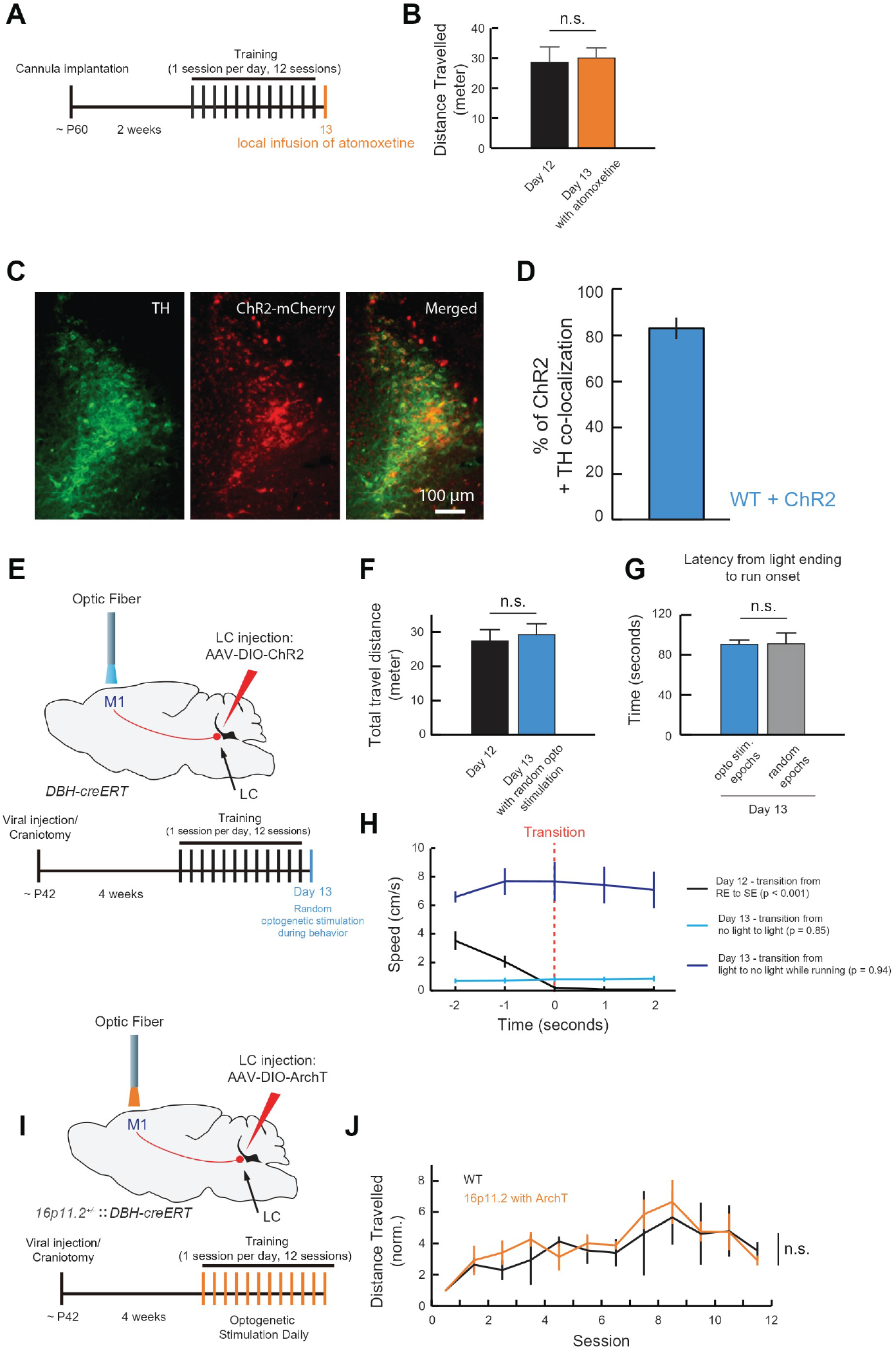
Closed-loop optogenetic validation and rescue experiments. **(A)** Experimental timeline. Mice underwent behavioral training with daily saline infusions for 12 days, followed by a single atomoxetine infusion on day 13 to assess whether atomoxetine administration affects movement expression. **(B)** Atomoxetine administration did not affect the general movement performance on the total distance travelled compared to the previous day (N = 3 mice). Wilcoxon rank sum test. **(C)** Representative images showing ChR2-mCherry (red), TH immunofluorescence (green), and co-localization (merge). **(D)** Mean fraction of ChR2-expressing cells co-localized with TH^+^ NA neurons (N = 5 mice). **(E)** Top, schematic of the virus injection and optogenetic manipulation. Bottom, experimental timeline with optogenetic manipulation in *DBH-creERT* mice. Mice underwent behavioral training for 12 days, followed by randomly delivered optogenetic pulses throughout the session on day 13 to assess whether optogenetic manipulation affected general motor performance. **(F)** Random optogenetic manipulation did not change the total travel distance compared to the session before (without optogenetic manipulation) (N = 3 mice). Wilcoxon rank sum test. **(G)** Random optogenetic manipulation did not produce any delayed effect on the subsequent RE initiation (N = 3 mice). The latency from the end of light stimulation to the onset of the next RE was calculated to assess the delayed effects on movement expression (from day 13), and it was compared to randomly selected epochs as a control (from day 12). Wilcoxon rank sum test. **(H)** Random optogenetic manipulation did not alter the behavior transition. Speed was calculated between behavior transition from Day 12 (RE to SE), Day 13 (no light to light), and Day 13 (light to no light during REs), and no changes in speed were observed in either light stimulation conditions. One-way ANOVA. **(I)** Top, schematic of virus injection and optogenetic manipulation. Bottom, experimental timeline with optogenetic manipulation in *16p11*.*2*^*+/−*^*::DBH-creERT* mice. Closed-loop optogenetic manipulations were applied daily throughout the entire training. **(J)** Optogenetic inhibition of excessive non-behavior rapid Ca^2+^ events rescues the delayed motor learning in the *16p11*.*2*^*+/−*^ mice. Results of the WT group are from the data in Fig7G.

## METHODS

### Animals

All animal experiments were conducted in compliance with the guidelines of the Canadian Council on Animal Care and received approval from the University of Ottawa Animal Care Committee. The colony of *16p11*.*2*^*+/−*^ mice was established through the mating of B6129SF1/J females (Jackson Laboratory, Stock No. 101043) with *16p11*.*2*^*+/−*^ males (Jackson Laboratory, Stock No. 013128). Subsequent mating involved continuously pairing male *16p11*.*2*^*+/−*^ descendants with B6129SF1/J females to enhance the colony size and reduce genetic variation. The *16p11*.*2*^*+/−*^*:DBH-CreERT* mice were generated from breeding *16p11*.*2*^*+/−*^ males with *DBH-CreERT* females (gift from Dr. Ellis Cooper, McGill University), characterized by the CD1 background. All mice were bred and accommodated in a facility with temperature range of 21−23°C and humidity maintained at 40−60% and have free access to food and water. The experimental mice were group-housed in plastic cages within a room where the light cycle was reversed (12 h/12 h). For the experiments, only water-restricted male mice within the age range of 8 to 14 weeks were utilized. Controls, unless explicitly stated otherwise, consisted of littermates sharing the same genetic background.

### Surgery for two-photon imaging

Mice were anesthetized with 1–2% isoflurane, and subcutaneous administrations of Baytril (10 mg kg–1) and buprenorphine (0.1 mg kg−1) were given to minimize infection and provide analgesia. An incision was made to extract a circular section of the scalp. For sensor imaging, viral injections (AAV-hSyn-NElm and AAV-hSyn-NEmut from BrainTVA; nLightG: gift from Dr. Tommaso Patriarchi [University of Zurich]) with no dilution were injected into 5 equidistant (200µm) sites of right M1 (0.3 mm anterior, 1.5 mm lateral, 0.2 mm depth from pia, ∼40 nL per site) in WT or *16p11*.*2*^*+/−*^ mice. For LC-NA axonal Ca^2+^ imaging, a viral injection (AAV1-Syn-Flex-GCaMP8M-WPRE from Addgene) with 1:6 dilution in saline was performed in the right LC (5.4 mm posterior, 0.8 mm lateral, 3.6mm depth from bregma, ∼600nL) of *16p11*.*2*^*+/+*^::*DBH-CreERT or 16p11*.*2*^*+/−*^::*DBH-CreERT* mice using a stereotaxic instrument (Kopf). For the LC-NA axonal Ca^2+^ imaging with chemogenetic manipulation, *16p11*.*2*^*+/+*^::*DBH-CreERT* or *16p11*.*2*^*+/−*^::*DBH-CreERT* mice were bilaterally injected with a viral mixture of AAV1-Syn-Flex-GCaMP8M-WPRE (Addgene) and AAV2/9-hSYN-DIO-hM3D(Gq)-mCherry (Neurophotonic Center, University of Laval) into LC. For excitatory neuron imaging in M1, AAV1-CamKII-GCaMP6f-WPRE-SV40 (University of Pennsylvania Vector Core, diluted 1:4 in saline) was injected into the right M1 at five sites (∼20 n1 per site), in which each injection was delivered at a depth of ∼250 µm from the pia, with injection sites spaced 250–500 µm apart. Following the injection, a custom-designed head-plate was affixed to the skull using instant glue (Krazy Glue) and dental cement (Lang Dental). A craniotomy of about 2 mm in diameter over the right M1 was conducted and a custom-made imaging glass window was placed over M1 and secured in position with dental acrylic. After the surgical procedures, mice received buprenorphine (0.1 mg kg^−1^) and dexamethasone (2 mg kg^−1^). Tamoxifen (Sigma Aldrich, 100 mg kg^−1^, dissolved in corn oil) was given on days 1, 4, 7 and 10 after the surgery through intraperitoneal injection in the *DBH-creERT mice*. Mice recovered for 4-6 weeks before behavioral training and two-photon 1magmg.

### Head-fixed rotating-disk behavior

Three days prior to the training/imaging sessions, mice underwent daily habituation while being head-fixed inside a plastic holding tube for 5 minutes. Training sessions were conducted for 1 hour each session, 1 session per day for 12 days. The hardware, software, and running threshold used for the bi-directional rotating-disk task have been described in our previous paper^16^. In this task, there is no sensory feedback or reward given to the animal. The disk can rotate in either direction, and the friction is low, so it is easy to rotate in the opposite direction if the animal pushes. Hence, mice can move in both directions, and it is difficult for the animal to maintain a steady running position at the beginning. Running Epochs (REs) were defined as a running period when a mouse’s velocity exceeded the running threshold and lasted at least 3 seconds (described in Yin *et al*. 2021)^16^, and an RE can last on average 11±3.7 seconds long on session 12. Stationary Episodes (SEs) were characterized as periods of behavior that do not fall under the classification of REs. WT littermates served as controls. Experimenters were kept unaware of the manipulations and genotypes of the animals.

### Postural Analyses

DeepLabCut (DLC)^43^ was used to label and track body parts using a using a high-speed (∼100 frames per second), high-definition night vision camera (Webcamera_USB) mounted above the animals. Video was recorded throughout learning across training sessions. Next, 240 frames of recorded behavior (selected by k-means clustering - a default DLC setting to select a variety of frames and optimize network training) were manually labeled — identifying nape, thorax, and sacrum — and used to train a deep neural network which then labels and reports coordinates during behavior. We then quantified movement variability as described in Yin *et al*. 2021^16^. Briefly, we isolated corporeal coordinates during REs, computed the angles created at the nape by the two most extreme sacrum points, and calculated the relative variance of these RE angles to assess posture and gait modification from sessions 1, 4, 8, and 12.

To identify specific postures, we used k-means clustering (k=5; validated by silhouette scoring and visualized with t-SNE) on the labeled RE coordinates. k (number of clusters/postures) was selected via the elbow method (tested against k=3, 4, 5, 7,10). The elbow method is a way to achieve good clustering performance while optimally selecting the minimum number of clusters; this prevents excessive/redundant clustering. Next, we sum the number of running frames in each posture (cluster), divided by the total frames in all postures, and normalize it to session 1 to calculate posture frequency. To compare clusters, we calculate the 2D-correlation of postures using their cosine similarity. We then computed the dot product of two posture vectors, divided this by their magnitude, and averaged across each of the coordinates (entire posture). To compare running features (duration, count) to postural features (correlation to primary posture, relative variance, postural entropy/uncertainty), we z-scored these values to allow a more direct comparison, then computed the correlation of these features’ trends across sessions 1, 4, 8, and 12.

### *In vivo* two-photon NA sensor imaging

The imaging sessions were conducted using a commercially available two-photon microscope (B-scope, Thorlabs) equipped with a ×16 objective (NIKON). Excitation was achieved at 925 nm (InSight X3, Spectra-Physics), operating at a frequency of 30 Hz. Two-photon images were recorded using ScanImage (Vidrio technologies), a software implemented in MATLAB 2015. For the NA sensor experiments, a single plane of images (512 × 512 pixels covering 400 × 350 µm^2^, ∼150-200µm from the pia) was continuously acquired while mice are engaged in running on the bi-directional rotating disk on Session 1, Session 7 and Session 12. For the non-learning control experiments, mice underwent training and imaging only on days 1 and 12 with no training or imaging in between. The same imaging plane was tracked across sessions for NA sensor imaging while different planes were used in axonal imaging in different sessions. Registration for NA sensor imaging was performed using ImageJ Turboreg Plugin implemented in MATLAB. Validation was performed using corr2() by computing pixel-wise correlation between 800 frames sampled at the beginning and at the end of a session and between different sessions across learning. These were compared within mice, and against randomly selected mice as controls. The experimenters were blinded to the genotypes of animals.

### NA sensor pre-processing

For NA sensor imaging GRAB_NE_, GRAB_NEmut_, and nLightG, motion correction was performed with full-frame cross-correlation image alignment (Turboreg plug-in ImageJ). A 7×7 ROI grid was used to segment the field of view (FOV) into 49 non-overlapping, equally sized ROIs, and the same FOV was followed throughout training. Florescent traces were extracted by averaging pixels within an ROI. Baseline fluorescence (F_0_) is calculated by taking the median of the raw trace, and a standardized fluorescence is calculated by subtracting *F*_0_ from the raw trace and dividing the result by the *F*_0_. The signal is passed through a Gaussian (kernel width 3sd) and median filter (medfilt1(), default settings) to remove speckle and salt & pepper noise, respectively. The filtered signal is then adjusted for drift using the MATLAB function ‘msbackadj()’, as in Feng et al 2019^17^, with i) a window size (1/12th the length of the fluorescence trace), ii) a step size of 90 using the spline regression method, iii) rlowess smoothing, and iv) a quantile value of 0.45. The filtered and baseline corrected traces were then z-scored to facilitate comparison across animals and days.

### NA sensor activity analysis

To determine NA levels from the fluorescent traces, we adapted an approach from the Harvey group^44^. For each ROI, z-scored traces from individual ROIs during SEs are concatenated across the entire session. Next, for each RE, a frame-matched epoch of the concatenated SE trace is randomly selected and compared to the RE trace. If 10 consecutive frames of the RE are higher than the frame-matched random SE selection, then the RE is considered elevated. The concatenated SE is then circularly permuted by a random number of frames to select another epoch with an equivalent number of frames, and the process is repeated. Following 1,000 repetitions, if the ROI yields 950+ elevated comparisons to the shuffled SE, then the ROI is considered active. For all the NA level analyses during REs, only active ROIs were analyzed. FOV was manually assessed for blood vessels and artifacts/viral expression, occluded ROIs were removed. To quantify the NA levels, we used an approach increasingly popular with neuromodulatory sensors. Rather than define discrete events, we computed the area under the curve (AUC) using the trapz() built-in MATLAB function. Re-activation index was computed as the percentage of REs that an ROI is active.

To categorize re-activation indices, we utilized the shape of their distributions to establish an activity threshold, along with an identity line (y=x) threshold that together would allow us to separate ROIs into increasing, decreasing, or stable categories. To compute a robust activity threshold, we computed the weighted average of the lowest local minimum (most conservative boundary) and the boundary derived by k-means clustering (k=2; increasing vs decreasing). ‘Stable ROIs’ had re-activation indices below threshold for both session 1 and 12, regardless of their position relative to the identity line. ‘Decreasing ROIs’ were defined as ROIs that exhibited re-activation index above the activity threshold in session 1 but below the identity line (y<x), indicating a decrease from session 1-12. ‘Increasing ROIs’ had an index above both the identity line, and above the activity threshold in session 12, indicating an increase from session 1-12. For spatial clustering composed of ‘patches’ of active ROIs, a binary ROI activity grid was created with the built-in MATLAB function bwlabel() using a connectivity of 4, such that connectivity between ROIs was limited to contact on the edges of the ROIs, excluding contacts on the corner; the size of the patches was attained using regionprops().

To validate the fidelity of GRAB_NE_ sensor, mice received craniotomy with AAV-hSyn-GRAB_NE_ injection in M1 as described above. Following recovery, mice were habituated to the two-photon image stage prior to imaging. On the imaging day, mice were first fully anesthetized using 1.5% isofluorane, imaged for 10 minutes before discontinuation of anesthesia. Imaging was done continuously for 30 minutes (10min anesthetized + 20min after discontinuation). Mice woke up within a few (1-2) minutes after the discontinuation of isofluorane, which was detected by the motion changes of the rotating disk. Awakening coincided with the maximal positive slope change of the fluorescence trace, and the awakening period was situated around this point. We quantified the change by using a window of 1,000 frames (10 bins, 100 frames each) around the awakening and compared to an equally sized window from the anesthesia period. The AUC was computed for each bin and averaged across bins for each period (awakening vs. anesthetized).

### Support Vector Machine for NA sensor imaging

A support vector machine (SVM) was trained for each mouse and session on four manually selected features of the NA fluorescence features: AUC, peak, mean, and# of active ROIs. These features were compiled by epoch (RE or SE), and 10-35% were held out for testing, similar to the canonical 20% hold-out. The model was provided balanced classes (same number of REs and SEs). Each model was assessed using the posterior probabilities fit resulting from the MATLAB built-in fitSVMPosterior() and tested using predict() to provide scores. SVMs use support vectors—or data points near the boundary as determined by training—to classify the data. Hinge loss is computed for each SVM using the loss() function. Scores were normalized and then pooled across mice within each session. True positive (TP), true negative (TN), false positive (FP),false negative (FN), and ROC curves are computed for each session using perfcurve(). Accuracy is computed by: (TP+TN)/(TP+TN+FP+FN). Individual TP, TN, FP, FN for one example mouse are represented in confusion matrices for session 1. Sensitivity, or true positive rate, is computed as: TP/(TP+FN). The distribution of accuracies for each session were compared using a T-test.

### LC-NA Axonal Ca^2+^ imaging

The same two-photon microscopy settings and imaging plane size used for NA sensor imaging were applied to axonal imaging. To avoid photobleaching in the axonal imaging, we imaged every 5min with 10min off throughout the 1 hour of training (4 x 5 min imaging epochs). Different planes were used in different imaging sessions. Following the acquisition of imaging data, lateral motion correction was implemented through full-frame cross-correlation image alignment (Turboreg plug-in in ImageJ). The MATLAB-based Suite2P program was employed for the automated identification of ROIs that represent individual LC-NA axonal fragments. The fluorescence signals associated with these ROIs were then extracted by Suite2P^45^. Gaussian filtering (sigma= 1) was applied to the raw traces of each ROI. The Matlab-embedded regression method msbackadj () was then utilized to compute the backbone trace for each ROI, which was used to calculate for persistent events. Subsequently, transient event-related traces were determined by subtracting the backbone traces from the filtered raw traces. To avoid treating individual axonal segments as a separate sample, axonal segments were clustered into neurons using principal component analysis (PCA) and hierarchical clustering on the z-scored persistent event-related traces from all the REs. In this calculation, principle components (PCs) that accounted for 80% or more of the variance were used for hierarchical clustering. The cluster number was determined by the highest score of the silhouette test. After clustering classification, each cluster was then defined as a functional LC-NA neuron. Once the clusters were determined, all the Gaussian-filtered raw traces of ROIs within each cluster were pooled together, and the average of these ROIs was used to represent the calcium traces of each functional LC-NA neuron. The persistent event-related trace of each LC-NA neuron was calculated using the regression method (msbackadj). Subsequently, the transient event-related trace for each LC-NA neuron was derived by subtracting the persistent event-related trace from the Gaussian-filtered raw traces of each LC-NA neuron. Linear trend was removed (detrend) from persistent event-related traces before further analysis. Following these definitions, a persistent event was defined as 60+ consecutive frames above 1 z-score, while a transient event was identified as 3+ consecutive frames above 1 z-score. Only ROIs with at least one persistent event during REs across the entire imaging session were included.

To validate the hierarchical clustering, the presence of persistent events during REs was assessed for each ROI throughout the imaging session. Each array was composed of binarized values for each RE in each ROI, where 1 indicated ‘presence of persistent event’ and 0 indicated ‘absence of persistent event’. We then calculated the variance of ROIs within an individual neuron and compared it to the variance of ROIs randomly selected within each RE. The median variance across all REs in an imaging session was then calculated. In addition, to compare the correlation of persistent events within individual neurons versus across neurons, binarized persistent event arrays of each ROI in the entire session were used to compute correlation values. These values were calculated both for ROIs within individual neurons and between different neurons. The median correlation value was calculated for each ROI.

To calculate the size of axonal patches with ‘abundant’ rapid events, we first determined a frequency threshold for high frequent rapid events using the 90th percentile of rapid event frequencies observed in WT mice. This threshold was subsequently applied to *16p11*.*2*^*+/−*^mice for comparative analysis. Following this, a distance threshold was then calculated based on the 10th percentile of the physical distances between axons exceeding the frequency threshold in WT mice. Axons within this defined distance threshold were classified as a single patch. The physical area of each identified patch were quantified using built-in convhull() function in Matlab.

To compare the spatial specificity between NA release and LC-NA axon activity, a 7×7 ROI grid was used to divide the LC-NA axonal imaging field into 49 non-overlapping, equally sized ROIs. Fluorescent traces were extracted by averaging the pixels within each ROI, followed by Gaussian filtering. Regression using the msbackadj() function were then applied to the traces of each ROI. Persistent and rapid events were identified using methods described above. Non-active ROIs, defined as those with no persistent events throughout the entire imaging session, were excluded from the analysis.

### Support Vector Machine-LC-NA axonal imaging

SVM was used to predict the occurrence of persistent events during REs. We utilized a window of 150 frames (equivalent to 5 seconds) leading up to the onset of each RE in each neuron for prediction. Four features, including rapid event frequency, mean, AUC, and average peak were computed and input into the SVM for prediction. A 5-fold cross-validation approach was applied, and the hinge loss function was used to calculate the loss for the classification ensemble model (loss). The accuracy was determined by subtracting the classification loss (including the false positive and false negative) for the cross-validated classification model (kfoldLoss) from 1.

### Pharmacological manipulations

Norepinephrine reuptake inhibitor, Atomoxetine, was locally infused into M1 of B6129S mice. Mice underwent the bilateral implantation of cannulae (PlasticsOne) into M1 (AP: 0.30 mm; ML: ±1.5 mm; DV: 0.25 mm) with adhesive cement (C&B Metabond, Parkell). After the implantation, dummy cannulae were inserted into the guide-cannula to prevent clogging. Mice recovered for 2-weeks before behavioral training. A 3-day habituation period was conducted as described above. To assess whether the atomoxetine treatment can result in delayed motor learning, mice received an infusion of atomoxetine (Millipore Sigma, 3 µg, 200 n1) into M1 20 minutes before each of the initial 3 training sessions, while saline-infused mice served as controls. To evaluate whether the atomoxetine treatment induces motor impairment, another cohort of mice received daily saline infusion throughout the 12-day training sessions, followed by a single dose of atomoxetine infusion (Millipore Sigma, 3 µg, 200 n1) on day 13.

Experimenters were blinded to animal groups. Local infusion was employed to enable the manipulation of NA signaling specifically within M1, without inducing global effects.

To investigate how atomoxetine administration changes NA release, mice underwent craniotomy with AAV-hSyn-NE1m injection. After weeks of recovery, mice were habituated to the disk apparatus and imaging rig. On the imaging session, we first imaged a ten-minute baseline period in a holding tube, followed by an intraperitoneal injection of atomoxetine. Controls were injected with saline. After 20 minutes to allow diffusion of the drug, mice underwent regular head-fixed rotating-disk training for 30 minutes with simultaneous two-photon imaging as described above. Intraperitoneal injection was employed as it was not feasible to perform two-photon imaging while administrating local infusion.

To investigate how atomoxetine administration alters M1 neuronal representations, mice received craniotomy with AAV1-CaMKII-GCaMP6f, and underwent a 3-day habituation period before head-fixed rotating-disk training and two-photon imaging. Since we cannot implant a cannula and have the cranial window at the same time, mice received the drug through intraperitoneal injection. On Day 1, mice received an intraperitoneal injection of atomoxetine (2 mg/kg) 20 minutes before training, and neuronal activity was recorded from hundreds of excitatory neurons in M1 through the craniotomy window. On Day 2, the procedure was repeated with an intraperitoneal injection of saline, and the same neurons were tracked from Day 1. Activity analysis, including the activity index and percentage of highly active cells, was performed as described in a previous publication (Yin et al., 2021). In brief, an individual neuron’s ΔF/F_0_ across all REs was averaged (Y) and compared with its own average ΔF/F_0_ during duration-matched SEs (X) in each session. Neurons spiked at least once throughout the imaging session were defined as active neurons and included in the analysis. Highly RE-active neurons were defined as those with a Y value greater than their corresponding X value, with a Y value exceeding a set threshold calculated by 50th percentile of the sorted ΔF/F_0_ values from all active neurons of all animals in the atomoxetine session. The activity index was calculated by the formula: *(Y* − *X)/(Y* + *X)*. Activity indices of all highly RE-active neurons were calculated, and the median of activity indices was calculated for each animal.

### Chemogenetic manipulations

To investigate how chemogenetic manipulation affects the LC-NA axonal activity, the surgery was described in the ‘Surgery for two-photon imaging’ section. Tamoxifen (Sigma Aldrich, 100 mg kg−1, dissolved in com oil) was administered via intraperitoneal injection on days 1, 4, 7, and 10 post-surgery. Following a 4–6 week recovery period, mice underwent a 3-day habituation period before head-fixed rotating-disk training and two-photon imaging. On Day 1, mice received an intraperitoneal injection of the hM3D(Gq) ligand JHU37160 (0.5 mg/kg) 20 minutes before training and imaging. On Day 2, the same procedure was repeated, but with an intraperitoneal injection of saline as control. Imaging and analysis of rapid and persistent axonal Ca^2+^ events were conducted as described earlier.

### Closed-loop optogenetic stimulation

A real-time closed-loop system was developed to automatically trigger LED activation in response to the mouse’s pupil size and running velocity. *16p11*.*2*^*+/+*^*_*. .*DBH-CreERT* (WT) mice underwent a viral injection (Neurophotonic Center, University of Laval, AAV2/9-EF1a-DIO-hChR2-(H134R)-mCherry, diluted 1:4 in saline) in the right LC (5.4 mm posterior, 0.8 mm lateral, 3.6 mm depth from bregma, ∼600 n1) using a stereotaxic instrument (Kopf). A head-plate was then affixed to the skull and craniotomy was performed to place a custom-made glass window over M1 and secured in position with dental acrylic. Mice recovered for 4 - 6 weeks before initiating the optogenetic manipulation experiments. DeepLabCut was initially trained with videos recorded by a night vision camera (Webcamera_USB), capturing changes in the mouse’s pupil size over time. During the behavioral manipulation experiments, the mouse’s pupil size was being tracked with the trained DeepLabCut, and simultaneously, the mouse’s running velocity was being recorded. Each training session contains two sections: the sampling period and the training period. In the sampling period, which consists of the first 5 minutes, only the mouse’s pupil size was recorded. The mean value of pupil size during this sampling period was then used to calculate the z-score in the following training period. During the training period (60 minutes as described above), z-scored values of the pupil size were calculated in real time. Arduino signals the DORIC LED driver (10-ms pulses, 10 Hz, ∼20 mW, 470 nm, Doric Lenses) to activate the diode only when the following two conditions are fullfilled: i) the z-score of the pupil size is smaller than 1 z-score, and ii) mice’s velocity is not exceeding the running threshold. *16p11*.*2*^*+/+*^. *:DBH-CreERT* littermates were injected with tdTomato (UPenn Vector Core, AAV1.CAG.Flex.tdTomato.WPRE.bGH, diluted 1:4 in saline) in the LC to serve as controls. Experimenters were kept blinded to animal groups. Script can be found in the following link: https://github.com/aaronjuma/MousePupilTracker/tree/main/docs)

To examine the correlation between running onset and pupil size dynamics, running speed was initially binarized using the running threshold. Subsequently, the Pearson correlation coefficient was computed between the changes in pupil size and binarized running speed across all REs. Randomly selected epochs of equivalent duration were utilized as a control to compute the correlation.

To assess whether optogenetic stimulations induce motor impairment, a separate cohort of mice underwent 12-day training sessions without optogenetic stimulation. On day 13, blue light pulses (10-ms duration, 10 Hz, ∼20 mW, 470 nm, Doric Lenses) were randomly delivered over the cranial window throughout the session (each stimulation is 12 seconds, 100x). Light stimulation timing and frequency were determined based on the closed-loop optogenetic manipulation experiment. To examine the immediate effect of optogenetic stimulation on the motor performance, speed was calculated from 2 seconds before and 3 seconds after the light stimulation onset. We then compare to the speed before and after REs with matching duration. To evaluate possible delayed effect of optogenetic stimulation on the motor performance, we calculated the shortest time interval between the end of light stimulation to the next RE, and we compared with randomly selected epochs as controls.

To further investigate whether inhibiting the non-behavior-related LC-NA activity restores the motor learning in the 16p11.2 deletion mice, another cohort underwent same craniotomy as described above but with viral injection of AAV2/9-CAG-Flex-ArchT-TdTomato in the LC bilaterally of *16p11*.*2*^*+/−*^*::DBH-CreERT* (16p11.2 deletion) mice. After a 4–6 week recovery period, mice underwent experiments with closed-loop optogenetic stimulation bilaterally. During the manipulation, pupil size was tracked using DeepLabCut, and running velocity was recorded as described earlier. An Arduino-controlled DORIC LED driver (10-ms pulses, 10 Hz, ∼10 mW, 590 nm, Doric Lenses) activated the diode under the following conditions: i) the pupil size z-score was smaller than 1 z-score, based on the 5 min pupil size recording described above, and ii) running velocity remained below running threshold. Optogenetic manipulations were applied in every training session.

### Statistical analysis

All data acquisition and analyses in this study were performed with experimenters blinded to the manipulations and genotypes of animals. Sample sizes were not predetermined using statistical methods but were consistent with those reported in previous publications^16,35,46,47^ and statistical power was tabulated and confirmed post hoc. The data collection process did not involve specific randomization procedures, but animals were randomly chosen to each experimental group based on their genotype. Statistical analyses were carried out using MATLAB. Unless otherwise indicated, all reported values are presented as mean ± standard error of the mean (s.e.m.). Statistical significance was determined by Student’s t-test, Wilcoxon rank-sum test, Kolmogorov-Smirnov test, z-test, permutation test, repeated measures one-way analysis of variance (rmANOVA) with Bonferroni corrections, or two-way ANOVA with post hoc Tukey’s test. The Wilcoxon rank-sum test was applied to data with a non-normal distribution at specific time points, while the Student’s t-test was used for normally distributed data. Data distribution was assessed using the Lilliefors test. All comparisons were conducted using two-tailed tests unless specified otherwise.

## Acknowledgements

We thank Y. Li for sharing the GRAB-NE1m sensor, B.H. Liu, J.C. Beique, V. Breton-Provencher, R. Naud, J. Beninger, and members of the Chen lab for comments and discussions, and B.H. Liu for providing feedback on the manuscript. This work was supported by grants from Canadian Institutes of Health Research (CIHR), the Scottish Rite Charitable Foundation, and the Brain & Behavior Research Foundation (NARSAD) for S.Chen, Canadian Neurodevelopmental Research Training Platform (CanNRT) Fellowship for X. Yin, and Autism Speaks Predoctoral Fellowships for N. Jones.

## Author Contributions

X. Yin, N. Jones, and S. Chen conceived the project. All the experiments were performed by X. Yin and N. Jones. X. Yin, N. Jones, and S. Chen analyzed the data and wrote the manuscript.

## Author Information

Correspondence and requests for materials should be addressed to S. Chen.

